# Guiding calcium carbonate formation *via* liquid phase separation of extremely charged coral protein AGARP

**DOI:** 10.1101/2024.06.04.597398

**Authors:** Barbara P. Klepka, Agnieszka Michaś, Tomasz Wojciechowski, Anna Niedzwiecka

## Abstract

Biomineralization *via* the non-classical crystallization pathway is postulated to involve a transient liquid phase of calcium carbonate formed in the presence of polymers. In the context of coral biocalcification, these polymers may include coral acid-rich proteins (CARPs) secreted into the skeletal organic matrix. However, direct evidence for the existence of this liquid phase containing proteins is lacking. Here, we show that the intrinsically disordered aspartic and glutamic acid-rich protein (AGARP), the first CARP cloned from the Great Barrier Reef scleractinian coral *Acropora millepora*, can significantly influence early stages of CaCO_3_ nucleation and crystal growth through liquid-liquid phase separation. We introduce the concept of a biologically relevant crystallization precursor: a liquid protein-calcium condensate composed of CARP molecules and Ca^2+^ ions, which forms as a result of liquid-liquid phase separation in a crowded environment. Our work bridges the gap between the liquid phase separation and biomineralization research.

## Introduction

Biomineralization mechanisms described within the framework of non-classical crystallization (NCC) theory involving putative liquid precursors are attracting increasing attention for their potential to revolutionize our understanding of crystal formation in biological systems^1^. Unlike the classical crystallization pathway, which assumes that crystals grow by direct atomic or ion attachment, NCC states that crystal formation proceeds through intermediate states such as prenucleation clusters (PNCs), amorphous precursors, or liquid phases^2^. The sequential emergence, coexistence, and transformation of these states, depending on thermodynamic conditions, remain subjects of ongoing debate. PNCs are thought to be thermodynamically stable and highly hydrated 1-3 nm clusters of ions^3,4^. Another transient state, amorphous calcium carbonate (ACC), has been found in a living biomineralizing organism – coral *Stylophora pistillata*^5^. The ACC entities were postulated to form within the tissue, then adhere to the surface of the skeleton and eventually crystallize into aragonite (CaCO_3_ polymorph)^5^. Additionally, the presence of polymers was suggested to induce a transient liquid phase, referred to as the polymer-induced liquid precursor (PILP)^6^. PILP droplets have been observed in the presence of polyacrylic acid by atomic force microscopy^7,8^, but there is no other direct experimental evidence of PILP to date. Conclusions about the hypothetical prior existence of a calcium carbonate PILP were inferred from structural properties of solid mineral phases at various stages of crystallization, rather than from direct observations of liquidity^9,10^. This has led to speculation that PILPs may actually be polymer-driven ACC cluster assemblies^11^. Resolving these uncertainties is critical to advancing our understanding of biomineralization processes and potentially aiding in the development of novel biomimetic materials.

In the context of coral skeleton formation, these polymers include lipids, polysaccharides, and notably, coral acid-rich proteins (CARPs)^12^ secreted into the extracellular matrix (ECM) at the interface between coral tissue and the aragonite skeleton^13^. To date, only four CARPs have been cloned and partially characterized^14^. CARPs from *S. pistillata* bind Ca^2+^ stoichiometrically and precipitate aragonite in seawater *in vitro*^14^. CARP3 has been shown to roughen CaCO_3_ crystal surfaces and influence polymorph selection based on the Mg^2+^:Ca^2+^ ratio^15^. Immunolocalization studies revealed that CARPs are localized near early mineralization zones, corresponding to calcium distribution patterns^16^. Additionally, CARP1 has been found to sequester and concentrate Ca^2+^ ions even in a non-biomineralizing organism *Nematostella vectensis*^17^. CARPs can also interact with other proteinaceous components of the skeletal organic matrix (SOM)^18^.

CARPs are characterized by low-complexity sequences rich in disorder-promoting glutamic acid residues^19^, which suggests that they can be classified as intrinsically disordered proteins (IDPs)^20^. IDPs lack stable three-dimensional structures and exhibit significant conformational flexibility, making them highly sensitive to environmental conditions^21^. Furthermore, they can undergo liquid-liquid phase separation (LLPS)^22,23^, forming biomolecular condensates. The capacity for LLPS^24^, flexibility and surface binding ability has been speculated to be crucial for calcium carbonate biomineralization^25^, although the exact biophysical mechanisms remain elusive.

Here, we report on a novel CARP, the aspartic and glutamic acid-rich protein (AGARP), recently identified in the stony coral *Acropora millepora*^26^, a model species, whose genome has been fully sequenced,^27^ for the ecological study of coral development,^28^ biocalcification,^29^ and climate change^30^. We provide experimental evidence that AGARP is an IDP that significantly affects both the early stages of CaCO_3_ nucleation and subsequent crystal growth *via* a liquid phase. We suggest the possible existence of a biologically relevant crystallization precursor, a liquid protein-calcium condensate (LPCC) composed of CARP molecules and associated Ca^2+^ ions. This LPCC is formed as a result of LLPS in a crowded environment that reflects the properties of coral extracellular SOM.

## Results

### The unique sequence of AGARP

AGARP^27^ is a highly conserved protein for the *Acroporidae* family (**Fig. S1**), which is the dominant group of reef builders. Surprisingly, except for its high Asp and Glu content (**Figs. 1a, k**), AGARP shows no sequence similarity to the known CARPs from *S. pistillata*^14^ (**Fig. S2**).

**Fig. 1.**
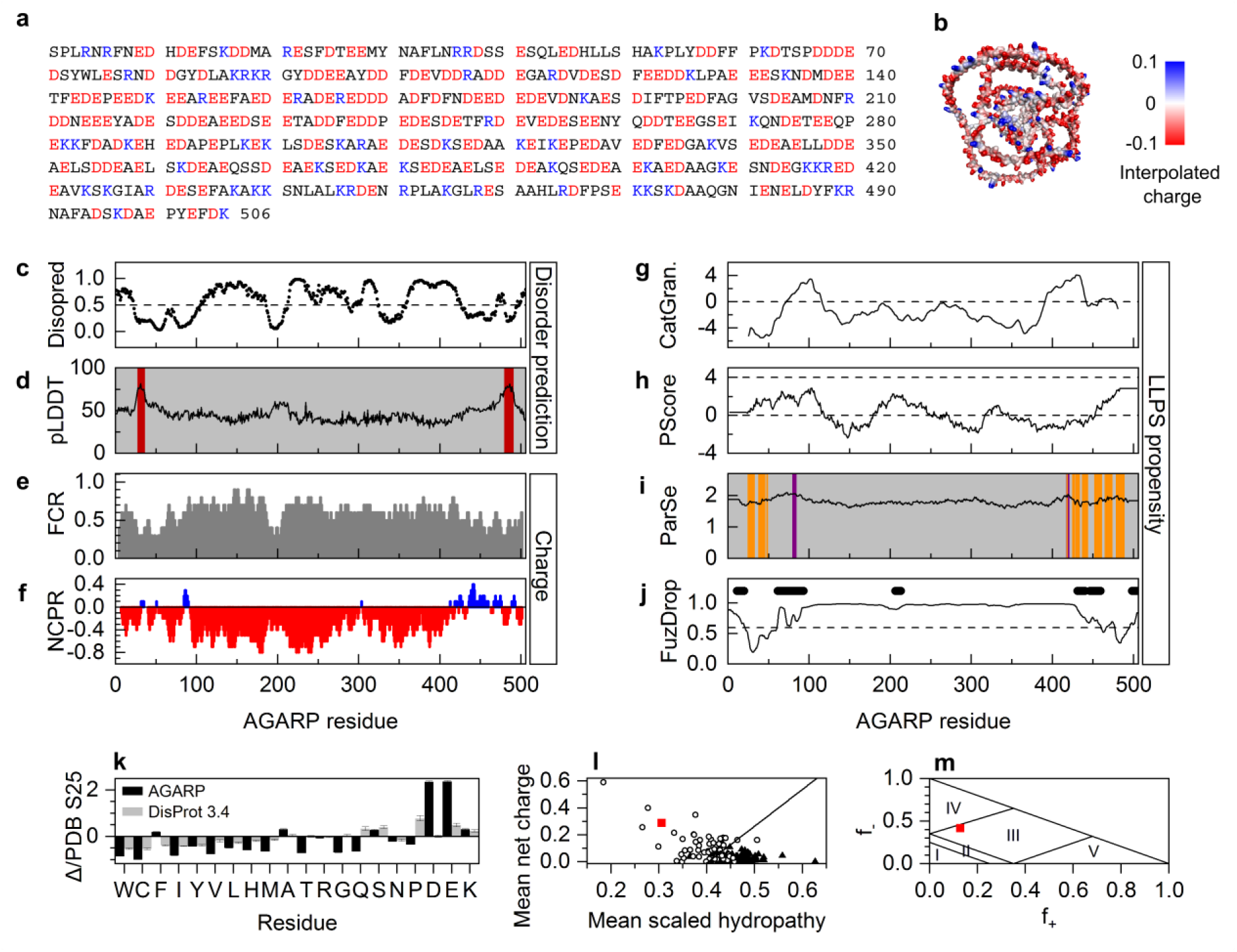
Predicted properties of AGARP. **a** AGARP sequence (Uniprot: B7W112) without signal peptide with marked acidic (red) and basic (blue) residues. **b** Predicted putative conformation shown as the solvent-accessible surface colored according to interpolated charge^31,32^. Bioinformatics analysis of AGARP; **c** disorder tendency^33^; **d** secondary structure prediction^31,32^, α-helices with pLDDT ≥ 70 marked in dark red; **e** fraction of charged residues distribution^34^; **f** positive (blue) and negative (red) net charge per residue; **g** – **j** LLPS propensity^35–38;^ **i** grey, intrinsically disordered regions not prone to LLPS; orange, regions that can fold to a stable conformation; purple, intrinsically disordered regions prone to LLPS; **j** thick black lines mark aggregation hot-spots; **k** composition profile^39^; **l** charge-hydropathy plot^40^ and **m** Das-Pappu phase diagram^34^, f_+_, f_-_, fraction of positively or negatively charged residues, respectively; III, strong polyampholytes; IV, negatively charged strong polyelectrolytes.

The conformation of AGARP is predicted to be almost completely disordered (**Figs. 1b, c**), with only local short-range α-helical propensity^33,31,41^ at the N- and C-termini (**Fig. 1d**), which correlates with the depleted fraction of charged residues (**Figs. 1e, f**). In the charge-hydropathy plot,^40^ AGARP ranks high in the subset of IDPs (**Fig. 1l**), but in contrast to most known IDPs, in addition to Glu residues, Asp and Phe are strongly overrepresented in AGARP, while the typical disorder-promoting Pro and Gln residues, as well as neutral Gly and Thr, are underrepresented (**Fig. 1k**). As a consequence, AGARP is an extremely charged protein with a net charge of –148 *e* per molecule and a linear charge density along the chain of −0.77 e/nm at pH 8.0, with an FCR exceeding 0.5 for most of the polypeptide chain (**Fig. 1e)**. The Das-Pappu phase diagram^34^ (**Fig. 1m**) identifies AGARP as a strong polyampholyte. However, the positive charges are clustered in short patches in the sequence (**Fig. 1f),** which is overall dominated by the negative charge. This suggests that AGARP may indeed exhibit the properties of a strong polyelectrolyte.

### Divergent predictions of AGARP’s propensity to phase separate

Four servers used to evaluate the LLPS propensity yielded puzzling results. catGranule^35^ identifies only two short LLPS-promoting regions (**Fig. 1g**) with a relatively low score for the whole protein, by 1.14 standard deviation (SD) larger than for the yeast proteome. The propensity predicted by PScore^36^ is 2.24 (**Fig. 1h**), which is below the confidence threshold of 4 SD. ParSe^37^ shows only negligibly short fragments at the N- and C-termini (**Fig. 1i**), with the score of 1.73 for the whole AGARP, below the 2.1 ± 0.2 limit for phase separating proteins.

The only server that predicts spontaneous LLPS of AGARP (*pLLPS* = 0.9945) is FuzDrop^38^ (**Fig. 1j**), which estimates the protein droplet propensity based on the conformational entropy of the protein chain, including not only the free state but also the binding entropy. In the case of CARPs, interactions leading to droplet formation can likely occur with proteins^26,42^, lipids, polysaccharides or counterions such as calcium or magnesium^43^ present in the coral SOM^44^. The LLPS propensity of AGARP should therefore be analyzed in the context of interactions with other SOM components, especially since the counterion binding may mediate the multivalent homotypic and heterotypic interactions between CARP molecules.

### Disorder and electrostatic collapse of AGARP

The intrinsic disorder of AGARP manifests itself in the hydrodynamic dimensions achieved by the protein due to the high like charge of the nearly polyelectrolytic molecule (experimental pI in the range of 3-4, **Fig. 2d**, calculated pI 3.9) and in the fluorescence spectrum with the emission maximum at ∼352 nm, indicating that the Trp indole ring is water exposed (**Fig. 2e**). The hydrodynamic radius, *R_H_*, of AGARP (and also of its His_6_-SUMO-AGARP fusion) is well above the 99 % CI of the linear Stokes-Einstein dependence for folded proteins approximated as diffusing hypothetical spheres of homogeneous density (*R_H_* = (0.788 ± 0.014)·*M*^1/3^; *M*, molecular mass; **Fig. 2c**). In terms of universal scaling laws, AGARP follows the power law for denatured proteins^45^, far from the characteristics of folded proteins determined under the conditions of this study (*R_H_* = (4.3 ± 0.2)·*N*^(0.312 ± 0.007)^) (*N*, number of residues in the protein chain; **Fig. 2c, inset**). The *R_H_*value of AGARP, 77 ± 5 Å (*M* = 58.3 kDa, *N* = 506, **Figs. 2a, c, S3, S4**) at 25 °C is larger than that of the HSA dimer, 44 ± 6 Å (133 kDa, *N* = 1170) and even larger than that of apoferritin, 58 ± 4 Å (480 kDa, *N* = 4200). This means that the AGARP molecule is described by an ensemble of expanded conformations close to a random coil. The hydrodynamic behavior of AGARP is temperature dependent, since the *R_H_* decreases to 66 ± 4 Å when the temperature is lowered to 10 °C (**Fig. 2b**).

**Fig. 2.**
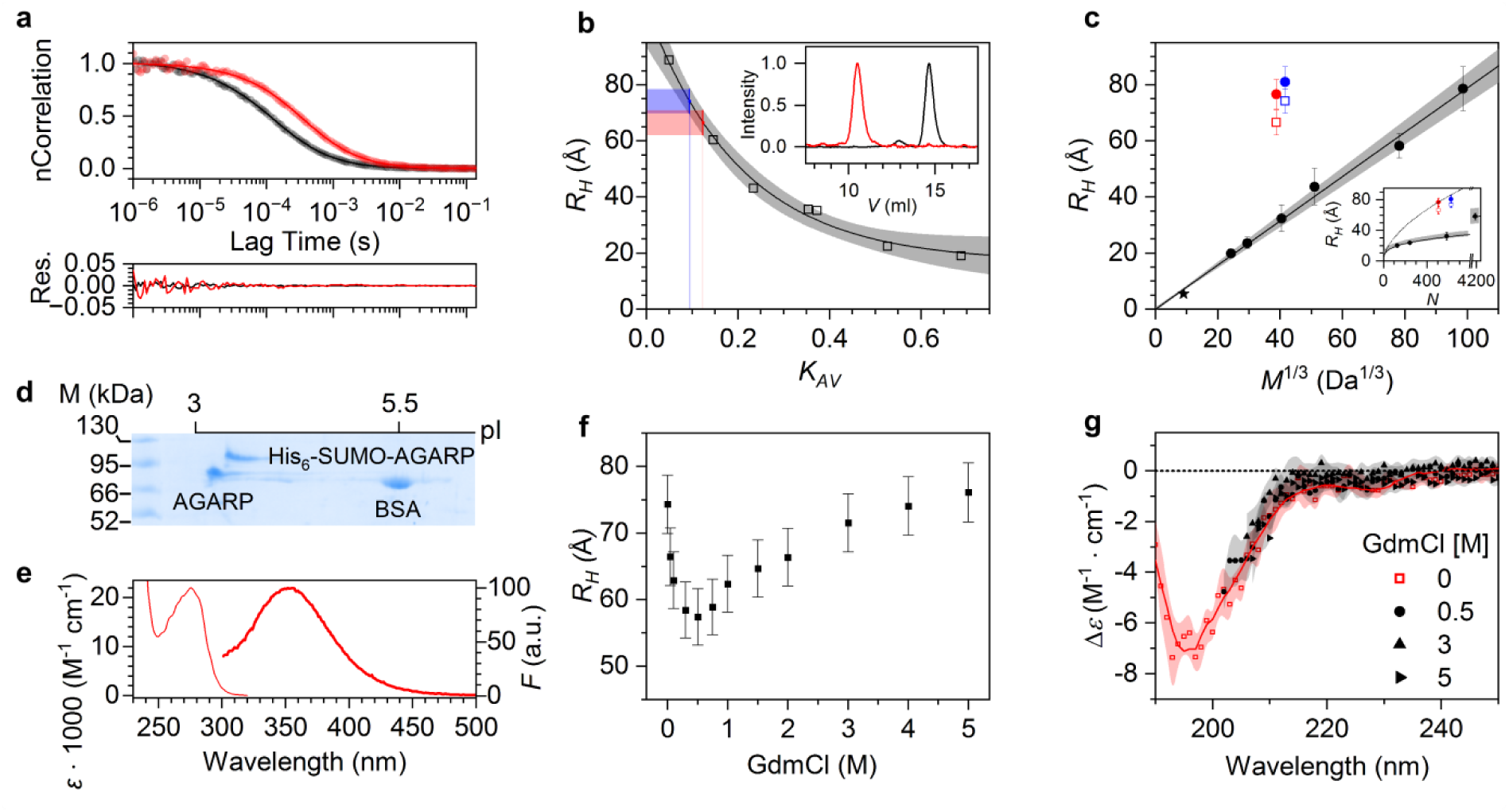
Experimental insights into biophysical properties of AGARP. **a** Normalized FCS autocorrelation curves (upper panel) for AGARP (red) and HSA monomer (black) at 25 °C with corresponding fitting residuals (bottom panel); dots, experimental points; lines, fitted models of diffusion. **b** Stokes radii, *R_H_*, of AGARP (red) and His_6_-SUMO-AGARP (blue) from SEC at 10 °C; (inset) SEC profiles of AGARP (red; 10.5 ml) and HSA (black; dimer at 12.9 ml, monomer at 14.7 ml); experimental uncertainty marked as shades colored correspondingly; SEC calibration function (black line, eq. 1) fitted to data points for model folded proteins (squares). **c** Dependence of *R_H_*on *M*^1/3^ and *N* (inset); AGARP (red circle) and His_6_-SUMO-AGARP (blue circle) at 25 °C and at 10 °C (hollow red and blue squares, respectively). Linear function fitted to FCS data for folded proteins at 25 °C (black circles) and free AF488 (star); (inset) power function of *R_H_ vs. N* (eq. 3) fitted to FCS data for folded proteins at 25 °C (solid line) and for denatured proteins (dotted line, *R_H_* = 2.21·*N* ^0.57^ Å)^45^. **b**, **c** Shaded gray areas, 99% confidence interval (CI) of the fits. **d** 2D IEF SDS PAGE of AGARP and His_6_-SUMO-AGARP; BSA shown for comparison. **e** UV/VIS absorption (thin line) and emission (thick line) spectrum of AGARP;. **f** Changes of AGARP *R_H_* with increasing GdmCl concentration as determined by SEC at 10 °C. **g** Normalized CD spectra of AGARP at different GdmCl concentrations at 10 °C; points, averaged experimental data; transparent shades, SDs colored correspondingly; solid line, fitted CD spectrum.

**Fig. 3.**
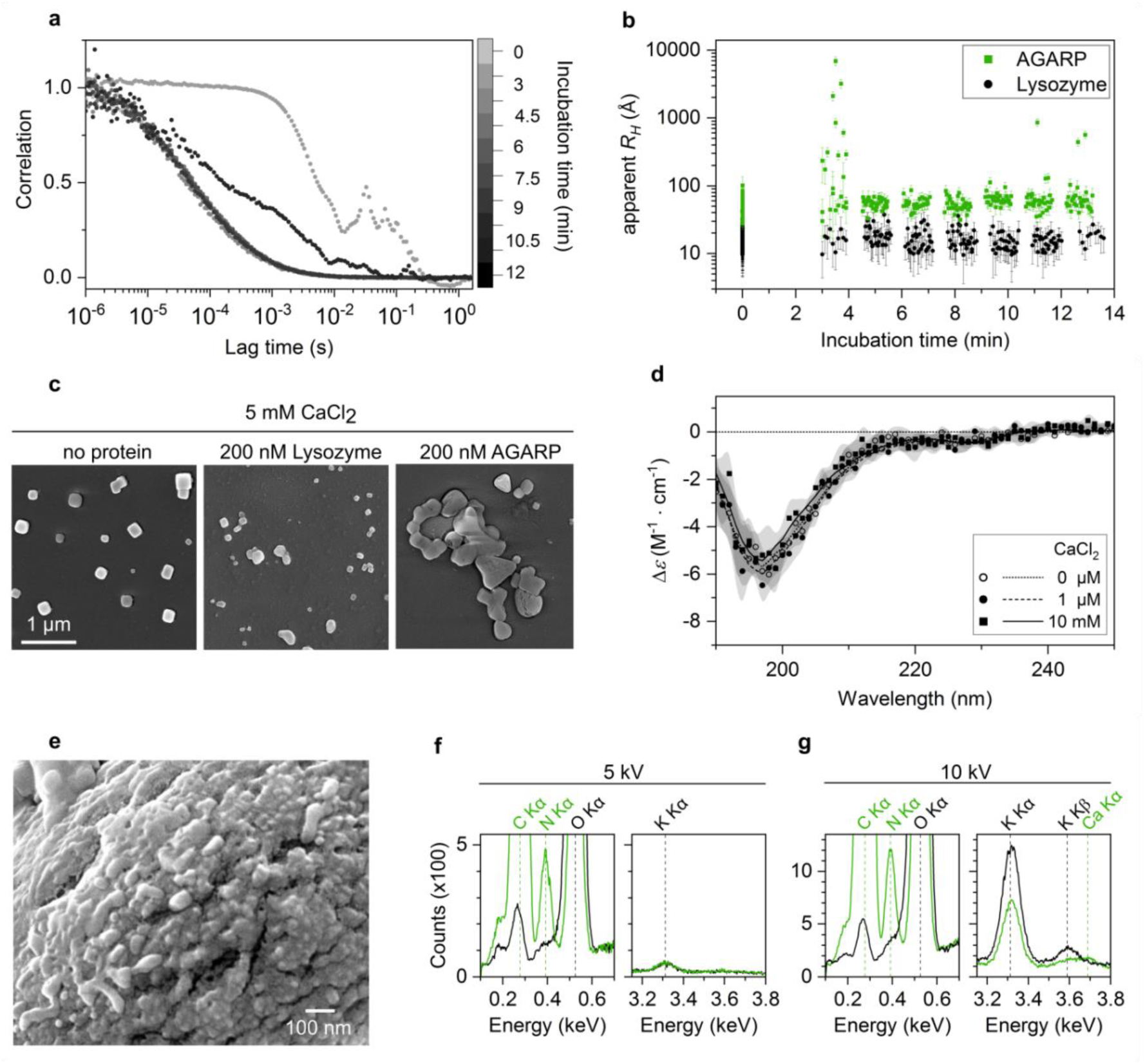
AGARP-driven formation of ACC. **a** Normalized FCS autocorrelation curves for AGARP (200 nM) at subsequent time points after the addition of 5 mM CaCl_2_; **b** Apparent *R_H_* values of AGARP-containing phases and lysozyme (control) shown as a function of incubation time with Ca^2+^; **c** SEM images of the phases deposited on the glass slide after the FCS experiment; scale bar is 1 μm in all images; **d** Normalized CD spectra of AGARP in the absence and presence of CaCl_2_; points, averaged experimental data; transparent shades, SDs colored correspondingly; solid lines, fitted CD spectra; **e** SEM image of the fragment of the phase formed after the FCS experiment analyzed by EDS; **f, g** EDS spectra obtained at (**f**) 5 kV and (**g**) 10 kV accelerating voltage; green spectrum, AGARP-containing CaCO_3_ phase; black spectrum, glass slide (control); dashed lines, K lines of elements.

**Fig. 4.**
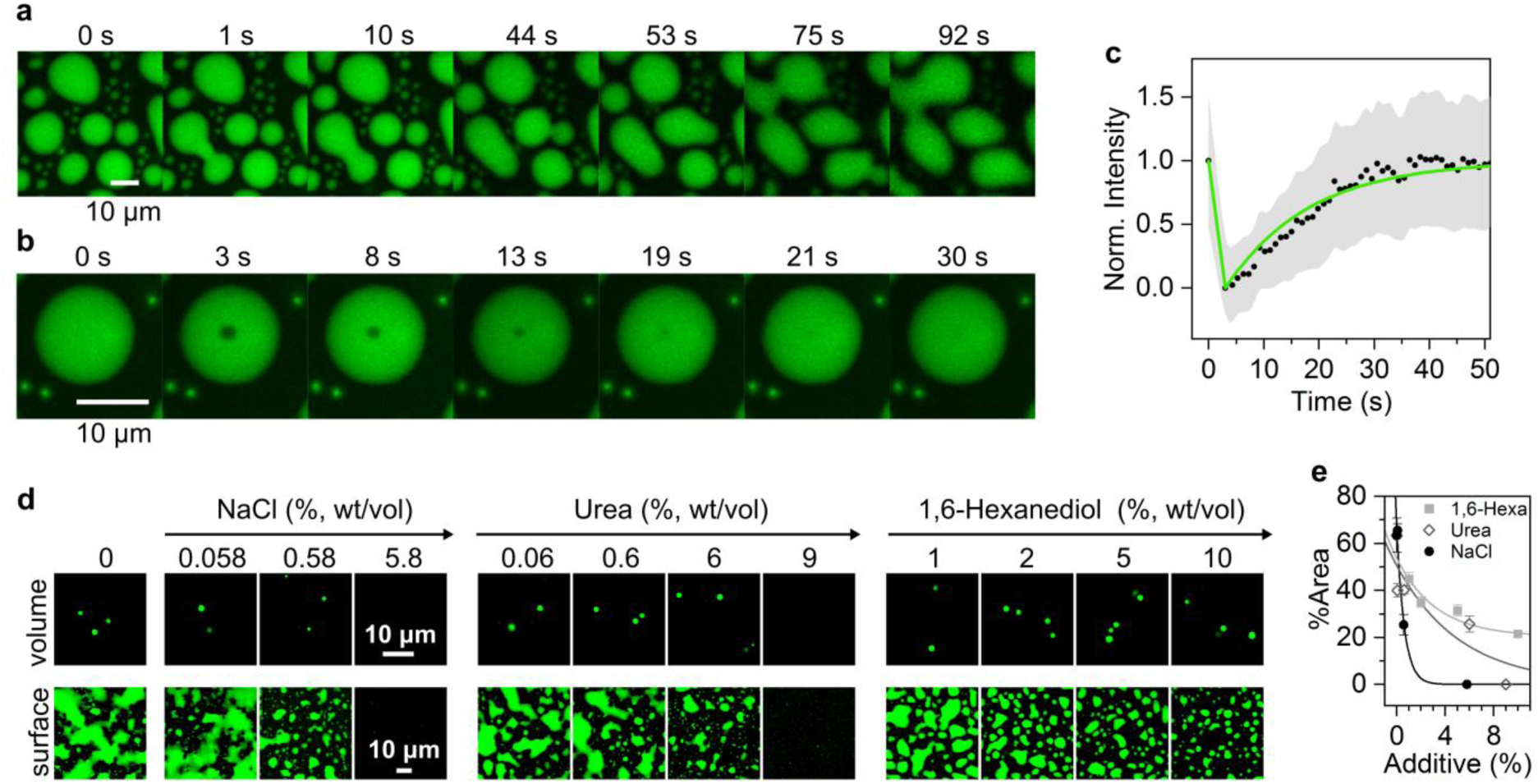
LLPS of AGARP-calcium condensates in crowded environment. Liquid droplets formed at 50 µM AGARP, 100 mM CaCl_2_ and 10% PEG 4k after ∼5 min: **a** Fluorescence confocal microscopy images of coalescence of AGARP-containing droplets; **b** Fluorescence recovery after photobleaching, and **c** the corresponding FRAP curve fit (green line) to measured intensities (black dots) with SDs (shaded area). Determination of the electrostatic mechanism of LLPS: **d** Fluorescence confocal microscopy images of droplets in the volume (upper panel) and liquid dense phase at the glass slide surface (bottom panel) formed at 5 µM AGARP, 50 mM CaCl_2_, 10% PEG 4k with increasing concentration of NaCl, urea and 1,6-HD; **e** corresponding quantitative analysis of the images; lines, decay function (eq. 5) fitted to experimental points (average ± SD from minimum 9 images). Scale bar is 10 μm in all images.

To gain an insight into the polyampholytic nature of AGARP, we measured *R_H_* in the presence of increasing concentrations of the ionic denaturant, guanidinium chloride (GdmCl). At lower GdmCl concentrations, the *R_H_* of AGARP decreases by ∼25%, reaching a minimum at 0.5 M, while at higher GdmCl concentrations, the *R_H_* gradually increases (**Fig. 2f**), similarly to what has been observed for other highly charged IDPs^46^. At low ionic strengths, the repulsion of negatively charged protein fragments is electrostatically screened by Gmd^+^ counterions, allowing the protein chain to relax. The subsequent expansion of the protein chain at higher ionic strengths could be attributed to the further non-specific counterion binding^46^.

Interestingly, these *R_H_* changes are not accompanied by secondary structure rearrangements as revealed by CD (**Fig. 2g**). A negligible fraction of α-helices and no long-range secondary structures in the form of parallel β-sheets were identified in AGARP both in the absence and presence of GdmCl. Other forms of local order, such as antiparallel β-sheets and β-turns, which are indistinguishable from random coil conformations by CD^47^, cover essentially 100% of the spectrum. The CD results show that AGARP exhibits features of an unstructured protein, even at the minimum of the *R_H_* value. This implies that the compaction of the AGARP chain under ionic strength in the 0.5 M range, comparable to extracellular conditions in invertebrates^48^, is an electrostatic collapse without folding or other disorder-to-order (*e.g.* fibrillation) transitions.

### Interactions of AGARP with Ca^2+^ ions

The biologically relevant counterion for CARPs is Ca^2+^, a building block of the coral CaCO_3_ skeleton. We therefore investigated the interactions of AGARP with Ca^2+^ using FCS and SEM. The *R_H_* of AGARP in calcium-free conditions is constant within 95% CI during a ten-minute FCS measurement (**Fig. S5**). The addition of Ca^2+^ to the protein solution shifts the individual autocorrelation curves of AGARP toward longer diffusion times (**Fig. 3a**) as a result of rapid protein aggregation due to charge inversion, which occurs in real time as AGARP molecules diffuse through the confocal volume. As shown in **Fig. 3b**, this shift corresponds to the formation of AGARP-containing aggregates with an estimated apparent *R_H_* of ∼100 Å to 1 μm, in contrast to lysozyme, for which no larger protein-containing particles are observed.

Under ambient CO_2_ conditions, AGARP triggers the formation of larger and clustered amorphous calcium carbonate (ACC) particles (**Fig. 3c**), which are deposited on the microscope slide after FCS measurements, in contrast to the 200-300 nm CaCO_3_ crystals formed in both the absence and presence of lysozyme, as revealed by SEM (see also **Fig. S6**). The surface of one of the largest AGARP-containing ACC aggregates (**Figs. 3e, S7**) exhibits noticeable roughness originating from ∼50 nm grains. The presence of AGARP in these particles was confirmed by EDS, since the peak corresponding to the Kα line of protein-derived nitrogen was detected at both 5 kV (**Figs. 3f, S8a**) and 10 kV accelerating voltage (**Figs. 3g, S8b**). The carbon Kα line, originating from both AGARP and CaCO_3_, was also detected at both AVs. Strikingly, the calcium Kα line was only visible after excitation with the higher energy electron beam. This suggests that AGARP is mainly localized in the outermost layer of the ACC, whereas Ca^2+^ tends to be located deeper. Monte Carlo simulations^49^ (**Fig. S9**) show that the AGARP-containing layer is at least ∼0.5 µm thick.

To address the question of how the AGARP conformation is affected by interaction with Ca^2+^ upon ACC formation, we measured CD spectra upon addition of 1 µM and 10 mM CaCl_2_ to the buffered solution and found that the protein backbone remains disordered (**Fig. 3d**).

### AGARP-driven liquid phase separation

To investigate the interactions of AGARP with Ca^2+^ ions under conditions that more closely mimic the presence of other biopolymers in the coral extracellular SOM, PEG 4k was used as a crowding agent. In contrast to the previously observed rapid precipitation of AGARP-containing ACC aggregates observed by FCS and SEM, under crowded conditions, AGARP solution undergoes binodal decomposition triggered by interactions with Ca^2+^ ions (**Figs. 4, 5**). The liquid nature of the emerging droplets is demonstrated by the observation of multiple coalescence events (**Figs. 4a, S10, Supplementary Movies 1, 2**) and the fluorescence recovery after photobleaching (FRAP) analysis (**Figs. 4b, c**) indicating a 100% mobile fraction and a fluorescence recovery half-time of 11 ± 2 s. As shown in the phase diagrams (**Fig. 5, S11**), the LLPS of AGARP is strongly dependent on the presence of the polymeric crowding agent, since at fixed protein and calcium concentrations the formation of the liquid dense phase occurs at higher PEG concentrations, whereas at lower PEG amounts only aggregates are detected. These observations suggest that molecular crowding is a prerequisite for the occurrence of the liquid phase.

**Fig. 5.**
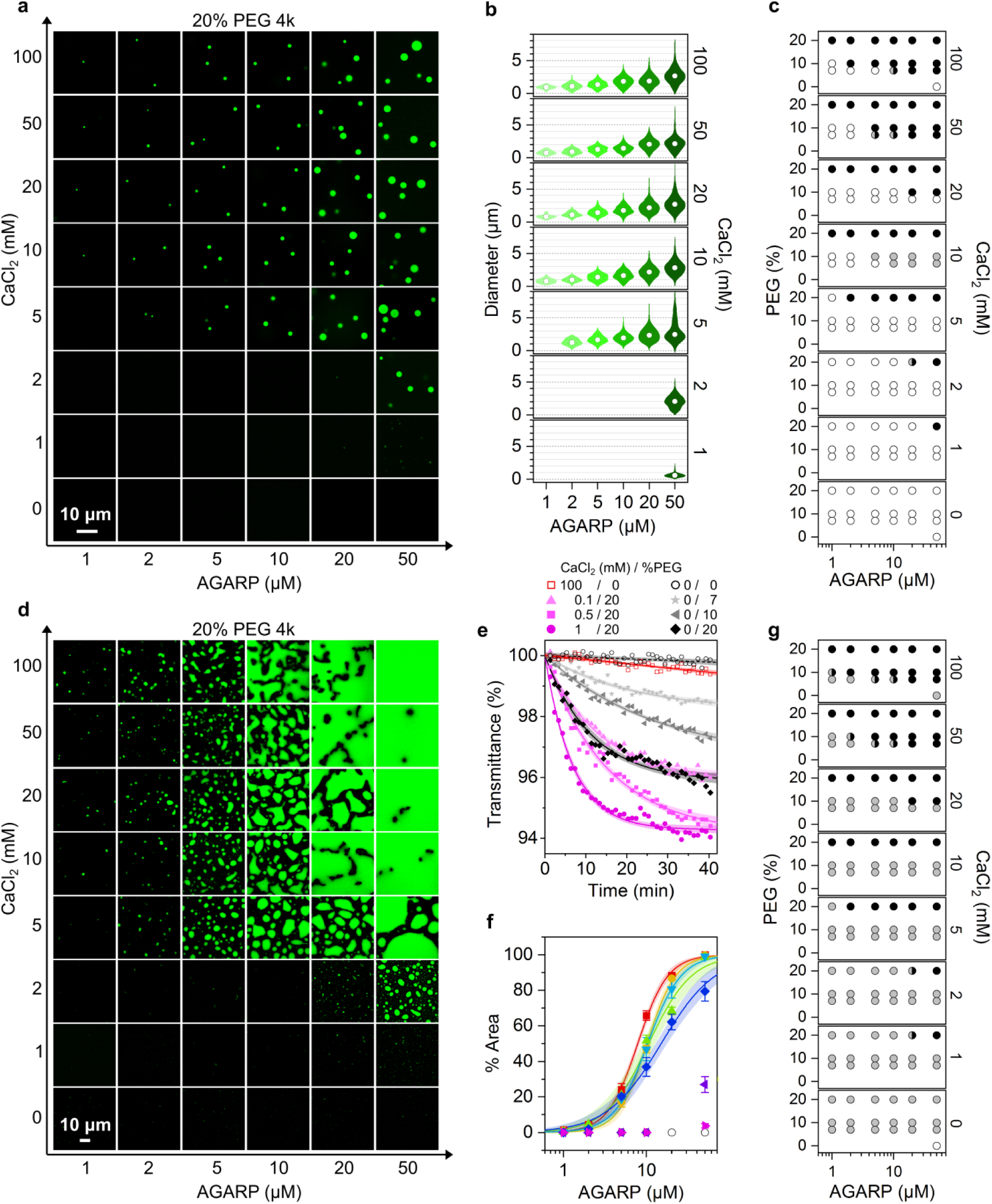
Quantitative analysis of AGARP-driven LLPS in crowded environment. **a** Fluorescence confocal microscopy images of AGARP-containing droplets formed at equilibrium in the volume part at increasing concentrations of AGARP and CaCl_2_, with PEG 4k at 20%. **b** Droplet diameter distribution as a function of AGARP and CaCl_2_ concentrations at 20% PEG; increasing green intensity corresponds to AGARP concentration (*N* from 25 to 2328 per distribution). **c** Phase diagram of LLPS in the volume part (5 μm above the slide); black, droplets; white, no dense phase; gray, aggregates; gray/black, both droplets and aggregates. **d** Fluorescence confocal microscopy images of the AGARP-rich phase accumulated at equilibrium on the slide surface at increasing concentrations of AGARP and CaCl_2_, with PEG 4k at 20%. **e** Turbidity analysis for AGARP at 50 µM; increasing intensity of magenta and grey indicate increasing concentration of CaCl_2_ and PEG 4k, respectively; lines, fitted function (eq. 6); shaded area, 99% CI. **f** Percentage of area occupied by the AGARP-rich phase at 20% PEG 4k as a function of AGARP concentration at different CaCl_2_ concentration (mM): 100 (red), 50 (orange), 20 (green), 10 (blue), 5 (navy blue), 2 (purple), 1 (magenta), 0 (black hollow); lines, fitted function (eq. 4) colored correspondingly; shaded area, 95% CI. **g** Phase diagram of LLPS observed at the slide surface; black, liquid dense phase; white, no dense phase; gray, aggregates; gray/black, both droplets and aggregates. Scale bar is 10 μm in all images.

The LLPS was analyzed 5 µm above and at the surface of the microscope slide (**Figs. 4**, **5**). Once the droplets appear in the bulk volume *via* the binodal decomposition, they coalesce rapidly and, when they are large enough, they settle by gravity (**Fig. S12, Supplementary Movie 3**) on the glass slide surface and undergo further multiple coalescence events. This leads to the establishment of an equilibrium of three liquid phases: one dilute bulk phase and two dense AGARP phases, the first being dispersed in the volume as fine droplets (**Fig. 5a**) and the second being the dominant phase on the slide surface (**Fig. 5d**).

To determine the mechanism of the LLPS, we performed the condensation inhibition experiments with additives that selectively interfere with non-covalent interactions, such as NaCl (weakening of electrostatic interactions and strengthening of hydrophobic contacts), urea (disruption of hydrogen bonds and van der Waals interactions), and 1,6-hexanediol (1,6-HD) (disruption of hydrophobic contacts) (**Fig. 4d**). The largest decrease in the number and diameter of droplets in the volume, which occurred at the lowest additive concentration, was observed for NaCl (**Fig. 4e, S13**). The diameter of the droplets in the presence of urea and 1,6-HD and the number of droplets formed with 1,6-HD remained unchanged. However, at the surface of the glass slide, the area percentage (%*Area*) of the dense liquid phase decreases exponentially for all additives. The largest relative %*Area* change and the lowest half inhibition concentration (c_1/2_) were observed for NaCl (**Fig. S14**), indicating the dominant electrostatic contribution to the LLPS mechanism involving AGARP.

The critical factors for the LLPS in the crowded environment that could be inferred from the confocal images are the concurrent presence of AGARP and calcium ions, since the absence of either of them seems to lead to the disappearance of the dense phase, as can be seen in the phase diagrams (**Fig. 5**) both in the volume part (**Figs. 5a-c**) and at the slide surface (**Figs. 5d, f, g**). Surprisingly, the distribution of droplet diameter depends exclusively on AGARP and is independent of the Ca^2+^ concentration (**Fig. 5b, S15, S16**), *e.g.*, for the maximum PEG concentration and regardless of CaCl_2_ concentration ranging from 2 to 100 mM, a threefold increase in the median of the droplet diameter is observed with increasing AGARP concentration from ∼0.8 μm at 1 μM to ∼2.5 μm at 50 μM. Conversely, the median diameter remains constant with rising Ca^2+^ concentration at fixed AGARP and PEG amounts, and with increasing PEG concentration (where the droplets are seen) at constant AGARP and Ca^2+^ concentration. Therefore, it is only the concentration of AGARP that determines the diameter of the droplet. Similar dependence on AGARP concentration was also observed for the number of droplets per mm^2^ (**Fig. S17**).

On the glass slide, the AGARP-rich phase occupies more space as compared to the volume phase (**Fig. 5d**). At lower AGARP and CaCl_2_ concentrations separated droplets can be observed, while at higher concentrations, the pattern observed for the dense phase is the result of multiple coalescence of droplets that have fallen by gravity onto a microscope slide from the volume phase and accumulate during the 2-3 hour equilibration period, although it could seemingly resemble a spinodal decomposition.

We investigated the changes of %*Area* of the AGARP-rich liquid phase accumulated on the glass slide surface after reaching the equilibrium with respect to the protein, Ca^2+^ and PEG concentrations. As shown in **Figs. 5f**, **S18**, the value of *%Area* increases sigmoidally with AGARP concentration according to the Hill equation (eq. 4), which suggests the cooperative nature of the process. This, in turn, is consistent with the fact that the analyzed pattern results from coalescence and accumulation. The quantitative analysis for different CaCl_2_ concentrations (**Fig. S19**) shows that the protein is the main quantitative determinant of the LLPS. The characteristic protein concentration describing the process is in the biologically relevant range of ∼10 μM and decreases only by 2-fold with the 20-fold increase in the Ca^2+^ concentration from 5 to 100 mM at 20% PEG 4k. An increase in the PEG concentration leads to a decrease in the characteristic concentration of AGARP, along with a simultaneous increase in the cooperativity coefficient, indicating the AGARP-specific driving force for the LLPS^50^.

The minimum concentration of CaCl_2_ sufficient to induce the formation of approximately micron-sized droplets detectable by confocal microscopy is 1 mM (**Figs. 5a, S20**). To evaluate the effect of micromolar calcium concentrations on the LLPS, we performed the turbidity assay based on the decrease in sample transmission (apparent increase in UV/Vis absorption) at 330 nm due to light scattering on a cloudy solution. The turbidity kinetics (**Figs. 5e, S21**) reveals that nano-sized droplets of AGARP are formed at 7, 10, and 20% PEG even in the absence of CaCl_2_, but the condensation is more efficient with increasing concentrations of CaCl_2_ in the sub-millimolar range. In an independent experiment, the spectra measured 10 min after the addition of calcium (**Fig. S22a**) show a bi-phasic process of the nano-droplets formation in the suspension (**Fig. S22b**). A small but statistically significant (P-value 0.00796) increase in A_330_ is observed below 0.8 mM (with a slope of 94 ± 15 μM^-1^), followed by a sudden transition above this concentration, consistent with the increase in the turbidity kinetic constant at 1 mM (**Figs. 5e, S21b**).

Both confocal images and turbidity measurements indicate that it is the presence of the CARP molecules that fuels the LLPS process under crowded conditions, while calcium ions play a supporting role.

### AGARP-controlled nucleation and growth of calcium carbonate

Having established that AGARP can stimulate the formation of either ACC clusters in simple water solutions or droplets under crowded conditions upon interactions with Ca^2+^, we investigated how these pre-crystallization states can tune the morphology of the emerging CaCO_3_ solid phases. The starting point for crystallization is the AGARP-Ca^2+^ complex in solution (**Fig. 6a**) or in liquid droplets (**Fig. 6h**) under PEG-free or 20% PEG 4k conditions, respectively. At all CO_3_^2-^ concentrations, AF488-labeled AGARP is highly incorporated into CaCO_3_, forming fluorescent structures (**Figs. 6b-g, j-n**). Interestingly, a complex dependence of the number of solid phase objects on AGARP and PEG concentrations was observed. With increasing CO_3_^2-^ concentration in the solution, CaCO_3_ deposits become smaller and more abundant (**Fig. 6b-d, i-k, o-q, t-v**), reflecting the typical stronger tendency of crystal nucleation at higher reactant saturation. However, under environmental CO_3_^2-^ conditions, solid forms (**Figs. 6b, e**) or droplets (**Fig. 6l**) are observed in the presence of AGARP, while no objects are formed without the protein (**Figs. 6o, t**). Moreover, in a CO_3_^2-^-enriched solution, crowding has a dramatic effect on nucleation in the presence of AGARP (**Figs. 6j, k** *vs*. **c, d**), while there is no effect on the amount of crystals in the absence of the protein (**Figs. 6u, v** *vs*. **p, q**). This implies that AGARP acts as a nucleator in cooperation with the crowding agent.

**Fig. 6.**
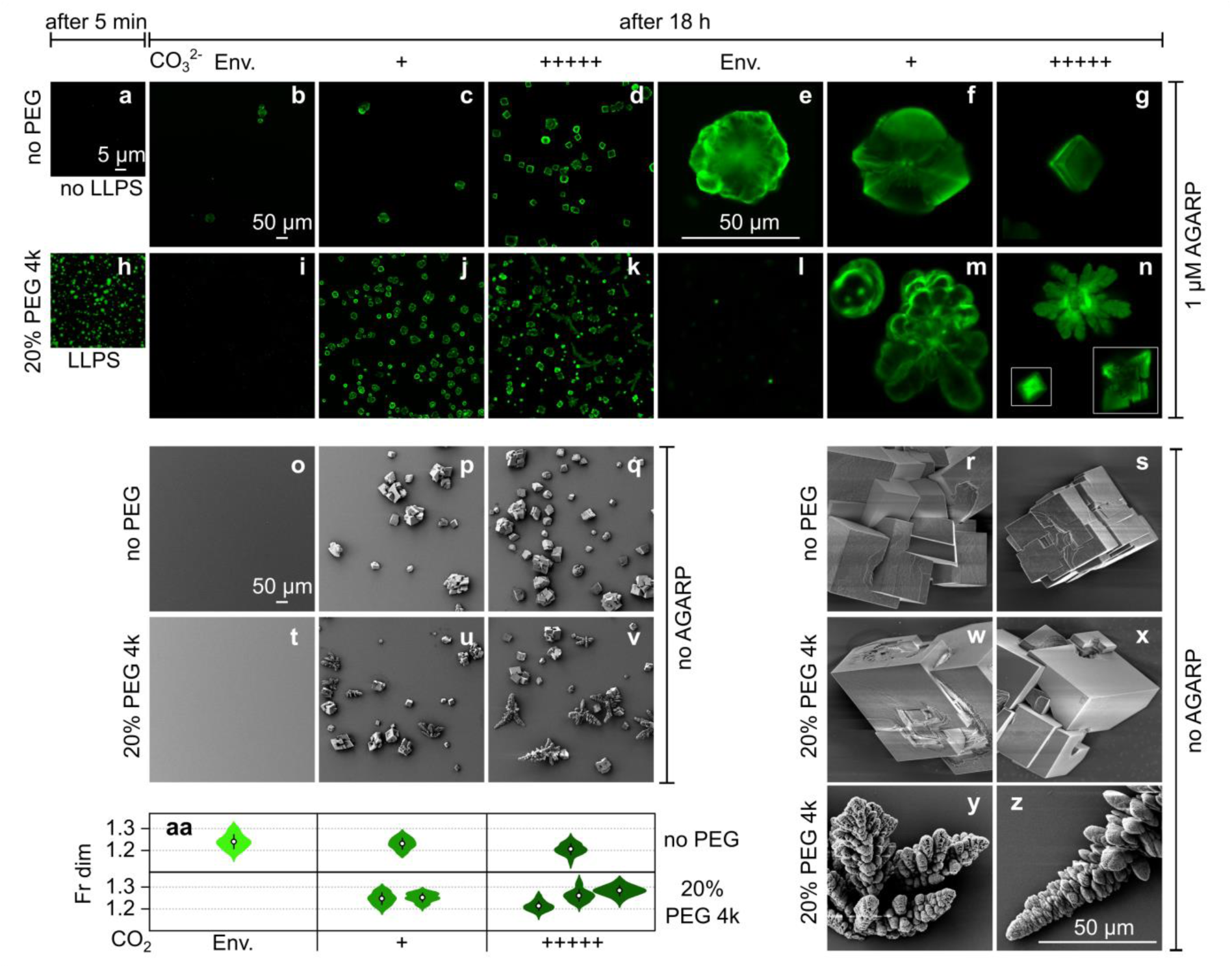
The growth of calcium carbonate is influenced by AGARP, CO_3_^2-^ availability, and crowding. **a, h** Fluorescence confocal microscopy images of phases formed on the microscope slide in the presence of 1 µM AGARP prior to the contact with CO ^2-^; **b**, **c**, **d**, **i**, **j**, **k** CaCO_3_ phases formed in the bulk (∼5 μm above the slide) at increasing concentrations of CO_3_^2-^ in the presence of AGARP and SEM images (**o**, **p**, **q**, **t**, **u**, **v**,) in the absence of AGARP, found in the 708.5 x 708.5 µm area of observation after 18 h crystallization; characteristic phases grown under the given conditions (**e, f, g, l, m, n** and **r, s, w, x, y, z**, respectively). Env., environmental source of CO_3_^2-^; +, 12.5 mg/cm^3^ and +++++, 62.5 mg/cm^3^ (NH_4_)_2_CO_3_ powder in the reaction chamber as a source of CO_3_^2-^ in solution, respectively. **aa** Fractal analysis of the phases shown in panels **e, f, g, m, n**. circle, mean; vertical line, SD. *N* from 43 to 86 per distribution. Scale bar is 50 μm in all images.

The CaCO_3_ phases grown with AGARP under the atmospheric or slightly enriched CO_3_^2-^ conditions with and without PEG have smoother and more rounded edges (**Figs. 6e, f, m**; only round droplets are present in **l**) compared to the conditions without protein (**Figs. 6r, w, y**). The effect of AGARP on changing the morphology of CaCO_3_ is most pronounced when the Ca^2+^ + CO_3_^2-^ ◊ CaCO_3_ reaction rate is slowed down by limited availability of CO_3_^2-^ both without and with crowding (**Figs. 6e** *vs*. **f**, **g** and **m** *vs*. **n**). However, the development of round edges under the influence of AGARP is more efficient for the crowded conditions where crystallization starts from LLPS (**Figs. 6m, n**) than from the solutions without PEG (**Figs. 6f, g**). Fractal analysis (**Fig. 6aa**) of the representative phases shown in **Figs. 6e-g, m-n** revealed that AGARP causes the fractal dimensions of CaCO_3_ phases to increase with the decrease in the amount of CO_3_^2-^, reflecting stronger edge development per unit area of the nascent crystals. In the absence of AGARP, no round edges are observed at all (**Figs. 6p-z**). Although fern leaf-like fractal structures of the calcium carbonate multicrystalline phase with strongly developed edges are formed under crowding conditions, they do not show any rounding (**Figs. 6y, z**).

### AGARP distribution in CaCO_3_ solid phases

The specific distribution of AGARP in the solid phases is strictly dependent on the reaction rate of CaCO_3_ formation. Under lower reaction rates, AGARP accumulates selectively in the center and the edges of the CaCO_3_ phases (**Figs. 7a, b, S23a**). This suggests that AGARP is responsible for nucleation and controls the edge development of the growing crystal. In addition, fibers with a cross-sectional diameter of 0.33 ± 0.08 µm are observed a few micrometers above the microscope slide (**Figs. 7a, b, S24**). The arrangement of these fibers, extending from the center of the crystal to the edges, likely reflects the protein’s mode of action: AGARP can initialize nucleation and then spread outward, modifying the crystal growth and edge formation.

**Fig. 7.**
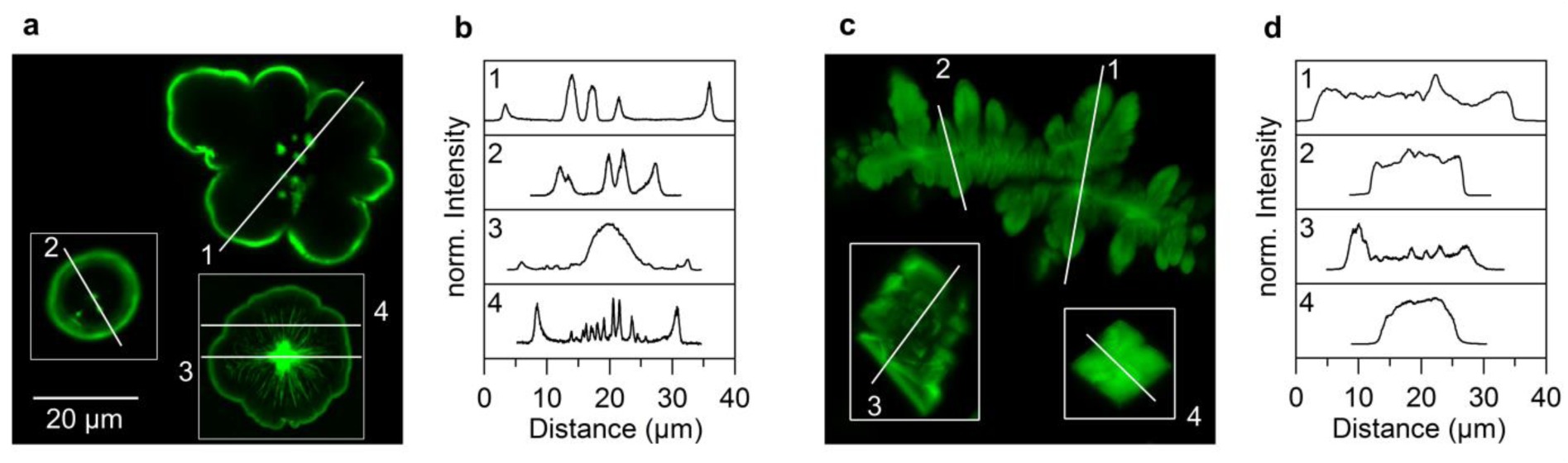
AGARP distribution in calcium carbonate depends on growth rate. **a** and **c** Fluorescence confocal microscopy images of phases grown in the CO_3_^2-^-enriched solution, + and +++++, respectively, as in Fig. 6; cross-section lines (white) shown in **b** and **d**, respectively. Scale bar is 20 μm in all images. AGARP at 1 μM, crowded conditions, 20% PEG 4k, 18 h of crystallization.

However, at a higher reaction rate of CaCO_3_ formation, AGARP is more evenly distributed within the phases (**Figs. 7c, d, S23b**), further confirming that the controlling role of proteins in determining morphology is limited under these conditions. Nevertheless, the slight increase in fluorescence intensity in the center of the CaCO_3_ phase in profile 1 (**Fig. 7d**) suggests that AGARP can still act as a nucleator even when the crystallization occurs more rapidly.

## Discussion

We have obtained and characterized the first CARP from *A. millepora*, AGARP, elucidating its physicochemical properties and its possible involvement in regulating the early stages of nucleation and controlling the growth of coral biomineral phases. Intrinsically disordered AGARP lacks a stable structure and has a hydrodynamic radius more than twice that of a folded protein of the same molar mass. AGARP first undergoes compaction and then swelling with increasing GdmCl concentration, similarly as other charged IDPs^46^. The high sensitivity of polyampholytes to the presence of salt^51^ is manifested in the interactions of AGARP with calcium. As a result of the electrostatic interactions with Ca^2+^, AGARP induces the emergence of highly clustered, amorphous CaCO_3_ particles, that can be attributed to ACC^52^, with AGARP accumulating predominantly on the outer surface. The structure of the AGARP-induced ACC clusters is similar to that formed with negatively charged dsDNA after 2 h incubation with stirring, which could influence the final structure^11^. With AGARP, however, ACC appears within minutes without agitation, suggesting that the effect of AGARP on ACC formation is stronger.

These results show that AGARP can be involved in the early stages of CaCO_3_ precipitation in the form of ACC, which is further supported by the findings that SOMs, including proteins, accumulate in the centers of coral calcification^53^ and that ACC was found to be one of the first precipitates formed in the pre-settled coral larva of *S. pistillata*^54^. The protein-driven stabilization of ACC was previously observed only for the mixture of proteins extracted from the coral SOM^55^. Here, we show that even a single CARP can stabilize ACC *in vitro*. The AGARP-induced formation of ACC appears to be a determinant of the crystallization rate, since it has been shown that the crystal growth occurring *via* ACC attachment is more than 100 times faster than *via* the classical crystallization pathway^5^.

*In vivo*, however, the crystallization processes in corals occur in the highly dense ECM, which is packed with proteins^18,26,56^, polysaccharides and lipids^44^. Under such crowded conditions, mimicked by PEG, AGARP undergoes LLPS, which is enhanced in the presence of Ca^2+^ ions. LLPS mediated by Ca^2+^ binding with the resultant local charge inversion has recently been shown for the N-terminal tail of the RNA-dependent DEAD box ATPases, Ddx4 and Ddx3^57^. The Ca^2+^-dependent LLPS process could be described in terms of the percolation theory^58^ where the AGARP molecules form biomolecular condensates due to homotypic multivalent electrostatic interactions between acidic residues, mediated by Ca^2+^ counterions acting as stickers.

This calcium-mediated LLPS model could be extended to include heterotypic interactions with other CARPs in the SOM. The critical effect of AGARP concentration on binodal liquid phase decomposition in crowded environment highlights the importance of the high multivalency of CARPs in regulating calcium carbonate nucleation and growth through both homotypic and heterotypic Ca^2+^-mediated interactions. This may provide a clue to explain how different coral species fine-tune their skeletal structure through the differential abundance of various CARPs, even though their sequences are not specific in terms of 3D structure and binding sites in the traditional sense.

The involvement of a liquid crystallization precursor, termed PILP^6^, has been previously described in the literature. However, the nature of PILP is qualitatively different from the liquid states observed in this work. It has been postulated that PILP is the liquid form of CaCO_3_ formed in the presence of a polymer^6^, and the idea focuses on the interaction of Ca^2+^ with CO_3_^2-^, as if their encounter complex was formed prior to polymer engagement. Here, the droplets are composed of AGARP molecules clustered by Ca^2+^ ions, which is more biologically relevant, since the secretory CARPs encounter Ca^2+^ ions earlier, during their biogenesis in the endoplasmic reticulum (ER), before reaching the extracellular environment rich in CO_3_^2-^ ions. Hence the CARP-Ca^2+^ complex formation in the ER should be considered as the first step in the CARP-mediated biomineralization. We thus introduce the term liquid protein-calcium condensate (LPCC) to describe the biomolecular condensate we observed as a biomineralization precursor.

The role of LPCC as a precursor of crystallization was confirmed by the crystallization experiments. We showed that AGARP tuned the morphology of the mineral phase when LPCCs were formed prior to contact with CO_3_ ^2-^ and crystallization proceeded slowly enough. AGARP was found to smooth and increase the development of the edges of the CaCO_3_ deposits. A less pronounced effect of AGARP in determining the mineral phase morphology was observed when the crystallization precursors were ACC particles.

The issue concerning the role of different types of proteins in CaCO_3_ crystal morphology modulation has been previously investigated (*eg*.^59,14,60,15^) and dates back to 1872^61^. However, the role of CARPs has only been studied in terms of polymorph selection and their ability to spontaneously precipitate aragonite^14^, which is strongly dependent on the concentration of Mg^2+^ ions^15^.

We propose a model for the non-classical crystallization pathway mediated by the export of CARPs in liquid condensates with calcium (**Fig. 8**). The AGARP protein, similarly as other CARPs^26^, has an N-terminal signal peptide (**Fig. S25**) that directs the protein onto the secretory pathway by targeting the nascent protein to the Sec translocon^62^ located in the ER membrane. The Sec translocon transports the translated protein to the ER lumen, a compartment where Ca^2^ ^+^ concentrations range from 100 μM to 1 mM^63^. We postulate that at this stage there is an interaction between CARPs and Ca^2+^ and their complex can be formed (1). From this stage on, CARPs are found in a highly crowded environment. The lumen of the Golgi apparatus and the secretory vesicles through which CARPs are transported to the plasma membrane, are packed with proteins and other biomolecules that increase the osmolality of the inner solution. Finally, the presence of various biopolymers incorporated into the skeleton^26,56^ suggests that the ECM at the tissue-skeleton interface is a highly crowded environment, although the exact composition is still unknown. In this work, we show that crowding triggers the formation of LPCCs (2), which further coalesce to form larger LPCCs (3). When the LPCCs are exposed to the environment rich in CO_3_^2-^, derived from the decomposition of HCO_3_^-^ produced from CO_2_ by carbonic anhydrases, predominantly secreted into the EM adjacent to the skeleton^26,64^, the crystallization process occurs (4). The emerging CaCO_3_ crystals have highly developed and smooth edges, similar to the native morphology of the skeleton^29,65^.

**Fig. 8.**
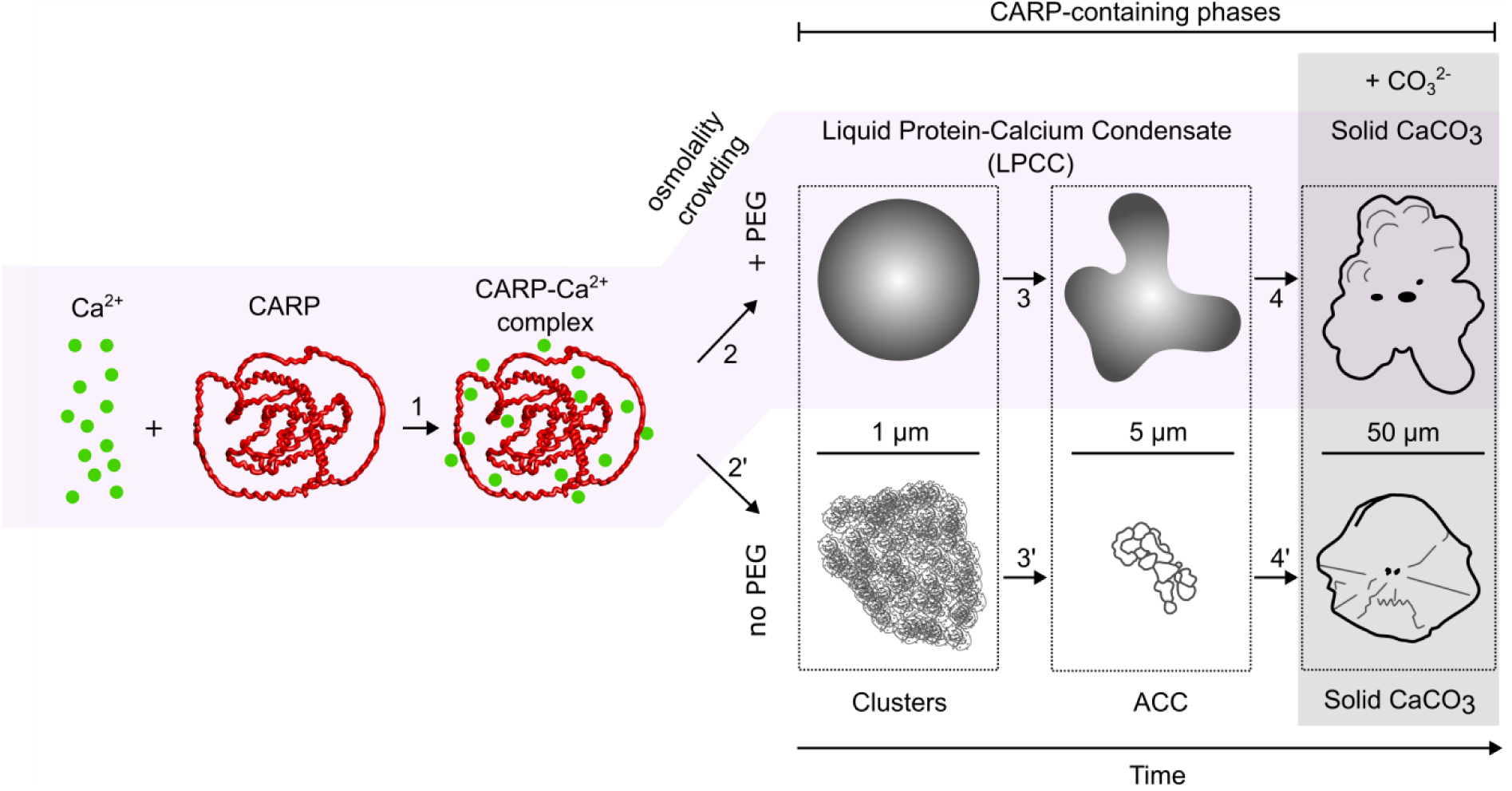
Model of the CARP-mediated non-classical crystallization pathway. When coral acid-rich proteins (CARPs) are exposed to the Ca^2+^-rich milieu in the ER, CARP-Ca^2+^ binding occurs (1). In highly crowded environment of SOM (violet area), LPCCs are formed *via* liquid-liquid phase separation (2). LPCCs coalesce (3) forming higher-order liquid phases. When LPCCs are found in the environment rich in CO_3_^2-^, crystallization occurs (4) and CaCO_3_ crystals with smooth and well developed edges appear. In less crowded conditions, CARP-Ca^2+^ complexes aggregate (2’) to form amorphous calcium carbonate (ACC) (3’). Once ACC is exposed to CO_3_ ^2-^, crystallization takes place (4’) and CaCO_3_ crystals with residual sharp edges are formed.

If the crystallization and precursor formation occurred in a low-crowded environment close to that of water (∼ 0.9 cp) in the absence of a polymeric crowder, the subsequent steps after the formation of the CARP-Ca^2+^ complex would be: (2’) aggregation of CARPs, driven by electrostatic interactions, (3’) formation of ACC and (4’) crystallization, leading to the formation of crystals with residual sharp edges. This pathway, however, would lead to the CaCO_3_ morphologies that could potentially injure the soft tissues.

Our results deepen the understanding of the involvement of the ACC-mediated crystallization pathway in the skeleton formation^5^. The formation of the CaCO_3_-based coral skeleton is an intricate process that depends not only on CARPs, but also on other biopolymers and environmental conditions. Most likely, different crystallization pathways intersect, one of which is mediated by a CARP-containing liquid precursor. However, a similar nucleation model has been proposed for mineral formation involving polystyrene sulfonate^66^.

CARPs share a common feature, *i.e.*, the negatively charged residues are overrepresented in their sequence^26,56^. However, the comparison of the results obtained for AGARP (**Figs. 6, 7**) and other proteins involved in biomineralization^60^ shows that the distribution of the proteins within the CaCO_3_ crystal, and thus their impact on nucleation and control of crystal growth, can be diverse. This in turn suggests that the existence of diverse CARPs is not limited to providing a high net negative charge, but that the composition of CARPs, which differ in the content and distribution of aromatic and hydrophobic residues in a given species, may be one of the factors determining subtle features of skeletal morphology.

The LPCCs have been discussed here as crystallization precursors. However, they can also perform two important functions: they can provide the SOM with a Ca^2+^ flux that is much stronger than Ca^2+^-ATPases-mediated ion exchange and free diffusion from seawater^44^, and they can facilitate the export of highly negatively charged CARPs across the membrane thanks to charge neutralization.

## Methods

### Reagents

All reagents were of analytical, spectroscopic or molecular biology grade: bromophenol blue (BPB), CaCl_2_·2H_2_O, CHAPS, glycerin, glycine, imidazole, NaOH, sodium dodecyl sulfate (SDS), thiourea, Triton X-100, urea, GdmCl (Carl Roth), dithiothreitol (DTT), NaH_2_PO_4_, β-mercaptoethanol (Sigma-Aldrich Merck), isopropyl β-d-1-thiogalactopyranoside (IPTG) (GeneDireX), Tris(hydroxymethyl)aminomethane (Tris) (Merck Millipore), polyethylene glycol (PEG) 4k (Fluka), HCl (Chempur), NaCl (Stanlab), ampicillin (BioShop), 96% ethanol (Pol-Aura). Proteins: chicken egg white lysozyme, bovine α-chymotrypsinogen A, bovine serum albumin (BSA) with monomer fraction ≥ 97%, human serum albumin (HSA), horse spleen apoferritin were purchased from Sigma-Aldrich Merck.

Buffers: A (50 mM Tris/HCl, 150 mM NaCl, 5 mM imidazole, pH 8.00), B (50 mM Tris/HCl, 150 mM NaCl, 500 mM imidazole, pH 8.00), C (50 mM Tris/HCl, 50 mM NaCl, pH 8.00), D (50 mM Tris/HCl, 1 M NaCl, pH 8.00), E (50 mM Tris/HCl, 150 mM NaCl, pH 8.00); F (25 mM Tris, 192 mM glycine, 0.1% SDS); G (50 mM Tris/H_2_SO_4_, 150 mM NaCl, pH 8.00); H (10 mM Tris/HCl, 100 mM NaCl, 5% glycerol, pH 8.00); I (50 mM NaH_2_PO_4_/Na_2_HPO_4_, pH 8.00); J (50 mM Tris/HCl, 150 mM NaCl, 0.5 mM EDTA, pH 8.00); equilibration buffer (6 M urea, 30% (w/v) glycerol, 2% (w/v) SDS in 0.05 M Tris/HCl buffer pH 8.8); rehydration buffer (7 M urea, 2 M thiourea, 4% (w/v) CHAPS, 40 mM DTT, 0.2% (w/v) Bio-Lyte 3/10 Ampholyte (BioRad), 1% bromophenol blue); Laemmli buffer (62.5 mM Tris/HCl pH 6,8, 1% (w/v) SDS, 10% (v/v) glycerol, 0,05% (w/v) bromophenol blue, 12.5% (v/v) β-mercaptoethanol). All buffers and AGARP samples used in experiments were treated with Chelex 100 resin (BioRad) to remove residual bivalent ions. pH of buffers with the addition of GdmCl or CaCl_2_ was adjusted individually.

### Protein expression and purification

DNA sequence corresponding to His_6_-tagged SUMO-AGARP (**Fig. S26**), purchased from Gene Universal, was cloned into pET-15b vector between NdeI and XhoI restriction sites (**Fig. S27**). Rosetta2(DE3)pLysS *E. coli* cells were transformed by heat shock with the prepared plasmid pET-15b. 100 μL of bacterial culture at a concentration of 10^4^ CFU/μg in LB Broth (BioMaxima) was incubated with 100-150 ng of the plasmid for 5 min on ice, 1 min at 42 °C and for 1 min on ice. LB Broth was added to the *E. coli* culture to a final volume of 1 mL and shaken for 1 hour at 37 °C. 100 μL of transformed cells were plated on LB agar with 100 μg/mL ampicillin and incubated O/N at 37 °C. A single colony was used to inoculate a pre-culture grown O/N in 10 mL LB Broth with 100 μg/mL ampicillin while shaking (250 rpm, SI-600R shaker, Lab Companion). Subsequently, 500 mL of the LB medium with 100 μg/mL ampicillin was inoculated with 5 mL of pre-culture and the cells were grown with shaking at 37 °C until OD_600_ ∼0.6. Then the expression of His_6_-SUMO-AGARP was induced by 0.2 mM IPTG. The protein was expressed at 15 °C for 4 h (**Fig. S28a**). The bacterial suspension was centrifuged and the pellet was stored O/N at -80 °C. The pellet was resuspended to the final volume of 20 mL in buffer A supplemented with supernuclease (Sino Biological) (250 U/1 g cell suspension), one tablet of protease inhibitors cOmplete Tablets, Mini, EDTA-free (Roche), and 2 mM DTT. The cell membranes were lysed by Triton X-100 added at a v/v ratio of 1:10 and sonication for 4 min on ice in cycles 30 s on, 30 s off at 50% of the maximal power in an ultrasonic homogenizer VCX130 (Sonics). Dissolved proteins were separated by centrifugation at 44,400×*g* for 30 min at 4 °C. The His_6_-SUMO-AGARP was purified on HisTrap HP (Cytvia) column by elution with increasing concentration of buffer B. The protein was subsequently incubated O/N with SUMO Protease (Kerafast) and SuperNuclease (SinoBiological) to cleave His_6_-SUMO tag and nucleic acids, respectively with simultaneous dialysis to buffer C. The tag was removed on WorkBeads NiMAC (Bio-Works) and nucleic acids on HiTrap Q FF anion exchange chromatography column (Cytiva) with increasing concentration of buffer D (**Fig. S28b**). The protein (**Fig. S29**) was further purified on the a Superdex 200 Increase 10/300 GL column (Sigma) in buffer E.

### Size Exclusion Chromatography (SEC)

SEC was performed on a Superdex 200 Increase 10/300 GL column (Sigma) at 10 °C with a flow rate of 0.5 mL/min, an injection volume of 100 μL or 500 μL, and the detection wavelengths of 215 nm and 280 nm. The column was equilibrated with the appropriate buffer before each experiment. For the determination of Stokes radii (*R_H_*) of AGARP forms, the setup was calibrated using standard proteins in buffer I: lysozyme (19 Å), α-chymotrypsinogen A (22.4 Å), BSA monomer (35.5 Å) and dimer (43 Å), HSA monomer (35.1 Å), and apoferritin monomer (60.3 Å) and dimer (88.8 Å). For the studies of interactions with GdmCl, AGARP was incubated on ice in buffer H with a given concentration of GdmCl for 30 min prior to measurements, and the SEC calibration was carried out in buffer H without GdmCl. Bigaussian function was fitted to AGARP (R^2^ = 0.963). The exponential function was fitted to calibration data points: *R_H_* = (*Y_0_* – Plateau)·e^−*A*·*K*^*_AV_* + Plateau (eq. 1), where: *Y_0_*, Plateau; *A*, fitting parameters; *K_AV_*, partition coefficient from SEC.

### Protein labelling

The proteins were labelled with AF488 NHS ester (Lumiprobe) according to manufacturer’s instructions in 100 mM phosphate buffer pH 8.45. The unbound dye was removed from proteins using SEC or Zeba Spin Desalting Columns 7K MWCO (Thermo Scientific).

### SDS–polyacrylamide gel electrophoresis (SDS-PAGE)

After expression and purification, samples were incubated with Laemmli buffer at 95 °C for 10 min. The SDS-PAGE was run in 12% polyacrylamide gel in buffer F at 120 V until the BPB left the gel. The gels were stained with Bio-Safe™ Coomassie Stain (BioRad).

### Isoelectric focusing (IEF) SDS-PAGE

Proteins His_6_-SUMO-AGARP, AGARP and BSA (50 μg each) were dialyzed against deionized water for 3 days (6 water changes) until there was no salt and the proteins precipitated. Immobilized pH gradient (IPG) (pH 3–10) strip (ReadyStrip, BioRad) was loaded with proteins dissolved in 125 μl of rehydration buffer and passively rehydrated O/N at 25 °C. 1 h after addition of the protein solution, the strips were covered with mineral oil (BioRad) to prevent evaporation. Isoelectric focusing was carried out in PROTEAN i12 IEF Cell (BioRad) at 20 °C at the maximum current of 50 μA/gel at 250 V for 15 min (rapid ramp), 4,000 V for 1 h (gradual ramp) and until 40,000 Vh (rapid ramp) was reached. The strips were later equilibrated for 10 min with 2.5 mL of equilibration buffer with 1% DTT (w/v) and afterwards with 4% (w/v) iodoacetamide. Before the SDS-PAGE separation, the strips were rinsed in buffer F and covered with an overlay solution composed of 0.5% (w/v) agarose in buffer F with BPB.

### Absorption and fluorescence spectroscopy

Absorption spectra were measured on Varian Cary 50 UV-Vis (Agilent Technologies) at a scan rate of 120 nm/min, 0.5 s integration time, 1 nm interval, for AGARP at 24.6 μM in buffer E; the buffer reference was subtracted.

Emission spectra were measured on Fluorolog 3.11 (Horiba Jobin Yvon) in a thermostated (20 ± 0.2 °C) semi-micro quartz cuvette with a magnetic stirrer, with path lengths for absorption and emission of 4 and 10 mm, respectively (119-004-10-40, Hellma), from 300 to 500 nm every 1 nm, with the excitation wavelength of 280 nm, 0.5 s integration time, excitation and emission slits of 1 and 10 nm, respectively, in the S/R mode, for AGARP at 0.8 μM in buffer E; the buffer spectrum was subtracted.

### Circular Dichroism (CD) Spectroscopy

A MOS-450/AF-CD spectrometer equipped with a Xe lamp was used to collect the signal from 190 to 260 nm, step 1 nm, 10 s integration time, in a thermostated quartz cuvette with the optical path length of 0.1 mm, in 4 replicates to exclude drift in time. The protein spectrum was then averaged and the averaged spectrum of the reference buffer was subtracted. CD spectra for AGARP at 7 μM in buffer H containing 0, 0.5, 3.0, 5.0 M GdmCl were measured at 10.0 ± 0.5 °C to compare with SEC. The measurements for AGARP at 8.13, 8.05, 8.03 μM in buffer G containing 0, 1 μM, 10 mM CaCl_2_, respectively, were performed at 20.0 ± 0.5 °C to compare with FCS. The final spectra were analyzed with BeStSel^67^.

### Laser scanning confocal microscopy imaging

Imaging was performed using an inverted laser scanning confocal microscope Axio Observer LSM 780 Zeiss, using MBS 488 beam splitter and BP 495-555 nm filter, excitation at 488 nm with Argon ion laser. LLPS was observed using Plan-Apochromat 63x/1.40 oil immersion lens. The phases of CaCO_3_ were visualized using C-Apochromat 40x/1.20 water immersion, air Plan-Apochromat 20x/0.8, or Plan-Apochromat 63x/1.40 oil immersion lenses.

### Fluorescence correlation spectroscopy

The FCS measurements were performed on the same confocal microscope equipped with C-Apochromat 40x/1.2W corr M27 water immersion lens and a ConfoCor3 kit, as described previously^68^. MBS 488 beam splitter and a BP 495-555 filter were used. Before each experiment, the samples were filtered through 0.22 μm membrane, and the 25 µL droplets were thermostated to RT of 25 °C for 3 minutes. The actual temperature inside the droplet was checked after every experiment with UT325 thermometer (UNI-T) equipped with a calibrated thermocouple with a diameter of 0.5 mm. The samples were excited at 488 nm with the Argon ion laser at 3% relative power. The setup: laser power, pinhole and objective ring, was calibrated on freely diffusing AF488 in deionized water^69^ and the structural parameter was determined separately for each microscope glass slide, pre-passivated with BSA. The triplet state relaxation time was measured in the independent experiment series individually for every protein at a higher relative laser power (15%). The triplet state lifetime and diffusion time of AF488 in buffer I for a given glass slide were fixed parameters in a two-component model of 3D diffusion that was fitted to experimental autocorrelation data to account for photophysics and a possible residual fraction of unbound dye. To determine *R_H_*, a single measurement lasted 3 s and was repeated 50 times in a series. The series were repeated for 3 or 4 independent droplets at 25.0 ± 0.5 °C. Spikes in the count rate related to the occurrence of oligomers or aggregates were excluded from further analysis. *R_H_* values were determined from the Stokes-Einstein equation taking into account the buffer viscosity and temperature in the droplet, as described previously^68^. The *R_H_* values were analyzed by a linear function, *R_H_*= *a·M*^1/3^ (eq. 2), where *M,* molecular mass, or by a power function from polymer diffusion theory^70^: *R_H_* = *R_0_·N*^ν^ (eq. 3), where *N*, the number of amino acid residues in the protein chain, *a*, *R_0_*, and the critical exponent ν, were fitting parameters.

Interactions of AGARP at 0.2 μM with Ca^2+^ ions were studied at 20 ± 0.5 °C in buffer J. Microscope glass slides were cleaned with 96% ethanol. Unpassivated microscope slides were used to avoid interactions of Ca^2+^ with BSA. Measurements before and after the addition of Ca^2+^ to the final concentration of 5 mM were run for 6 s, in 10 repetitions in a series. The series were repeated 7 times with a 30 s interval. The experiments were repeated in 3 independent droplets. Lysozyme was used as a reference.

### Scanning electron microscopy (SEM) and energy dispersive X-ray spectroscopy (EDS)

The phases that deposited on the microscope slide after FCS measurements with calcium and the phases grown during *in vitro* CaCO_3_ precipitation were then visualized using SEM. The elemental composition of the precipitates was determined using EDS. The samples were coated with a 5 nm layer of gold using a Q150T Sputter coater (Quorum Technologies) and inspected under Carl Zeiss Auriga CrossBeam SEM-FIB equipped with QUANTAX 400 Bruker EDS XFlash 7100. SEM images were collected at 5 kV accelerating voltage (AV) using an InLens SE or SESI detector for secondary electrons. EDS spectra we acquired at 5 and 10 kV AV.

### Coalescence and fluorescence recovery after photobleaching

To spot the early stages of droplet growth, liquidity and coalescence, the appropriate stock solution containing AF488-labelled AGARP and PEG 4k was placed on the silanized microscope slide and later CaCl_2_ stock solution was added to the final concentrations of 50 µM AGARP, 100 mM CaCl_2_ and 10% PEG 4k. Immediately thereafter, coalescence events were recorded by capturing images at 500 ms time intervals using the laser confocal microscope. After ∼5 min, three selected droplets were bleached with 100% laser power. The fluorescence recovery was observed for 52 s. The results were analyzed with FRAPAnalyzer 2.1.0 taking into account the contribution from background and bleaching during the measurement (references). A single exponential model was fitted, and the data were rescaled.

### Liquid-liquid phase separation experiments

Solution of AGARP labelled with AF488 was added to stock buffer E solutions containing appropriate amounts of PEG, CaCl_2_ and additives (NaCl, urea, 1,6-HD), for condensate dissolution experiment, and mixed by gentle pipetting in a test tube. The 4 µL mixture was placed on the unpassivated microscope slide with chambered cover glass (3 mm diameter, 1 mm depth, Grace Bio-Labs) and covered with another slide to prevent evaporation. The droplets or phases on the surface of the microscope slide and ∼5 µm above it were visualized after ∼3 h of incubation at 25 °C (4 to 22 images per condition and localization).

### Quantitative analysis of droplets and AGARP-containing phases

The percentage of area (*%Area*) occupied by the AGARP-containing phases on the surface of the microscope slide and the diameter of droplets ∼5 µm above the surface were determined from images using ImageJ. The Hill function: *%Area* = 100·*c^n^*/(*k^n^ + c^n^*) (eq. 4) turned out to be statistically the best (R^2^ from 0.976 to 0.999) for analyzing *%Area* as a function of protein concentration, *c*, where *n* is the apparent cooperativity coefficient, and *k* is a characteristic Ca^2+^ concentration at 50% of the observed process.

The exponential function *%Area* = *A*·exp(-*c*/*c_1/2_*) + *%A_0_* (eq. 5), where *A* is the relative area percentage change, *c_1/2_* is the half inhibition concentration and *%A_0_* is the plateau, was fitted to the *%Area vs*. NaCl, urea and 1,6-HD concentration data points; R^2^: 0.989, 0.786, and 0.944, respectively.

### Turbidity assay

The measurements were performed in two ways. AGARP solution was added to stock buffer E solutions containing appropriate amounts of PEG and CaCl_2_, (1) gently mixed twenty times in a test tube, or (2) mixed on a magnetic stirrer for 90 seconds and transferred to a 10 mm path length quartz cuvette. Absorption spectra from 250 to 700 nm were measured at 1-minute time intervals for (1) 40 or (2) 30 minutes at a scanning speed of 600 nm/min, 0.1 s integration time, every 1 nm, against a reference buffer with PEG and CaCl_2_. The starting point was (1) 1 min 20 s after mixing or (2) 2 min 10 s after the first encounter of two stock solutions. Absorbance was converted to transmittance and a one phase exponential function *%T* = 100+(*T_min_*-100)*(1-exp(-*K*\**t*)) (eq. 6) (R^2^ from 0.925 to 0.979), where *%T*, transmittance at 330 nm, normalized to the first data point; *T_min_* – minimal transmittance; *K* – rate constant; *t* – time; was fitted to time-dependent measurements using the (1) mixing method.

### *In vitro* calcium carbonate precipitation

The crystallization experiment was carried out for 18 h at 25 °C on a unpassivated microscope slide using a Press-to-Seal silicone isolator (9 mm diameter, 0.5 mm deep, Invitrogen) in a 25 µL droplet containing 100 mM CaCl_2_ without or with 1 µM AF488-labelled AGARP and without or with 20% PEG 4k in buffer E with either 0.4 mg (12.5 mg/cm^3^) or 2 mg (62.5 mg/cm^3^) of solid (NH_4_)_2_CO_3_ as a source of CO_3_ ^2-^. A CaCO_3_ precipitation was also performed in the atmospheric source of CO_3_ ^2-^, in this case the droplet was covered with a coverslip after 1 h of incubation. The CaCO_3_ particles grown in the presence of AF488-labeled AGARP were visualized by confocal microscopy and those grown in the absence of the protein were examined by SEM.

### Fractal analysis

ImageJ and FracLac plugin were used. The confocal images were rescaled to identical pixel dimensions and resized accordingly to obtain the same percentage of area occupied by the CaCO_3_ phases. The brightness was automatically adjusted, the threshold was set identically for all images to ensure visibility of the CaCO_3_ phase contours. The holes inside the phases were filled and the phase contour was outlined. The image quality-related artifacts was removed (**Fig. S30**). The fractal analysis was performed using 100 positions for grids, power series for calculating the series of box sizes and default number of box sizes and parameters for smallest and largest box size. Only fractal dimensions determined with r^2^ ≥ 0.999 were considered in further statistical analysis.

### Bioinformatics Analysis

Using AGARP (UniProtKB B7W112) without a signal peptide as a query sequence, the BLASTp algorithm was used to find homologous proteins. The masses and pIs of AGARP and His_6_-SUMO-AGARP were calculated using ProtParam. Disorder tendency was predicted by DISOPRED3^33^. Das-Pappu phase diagram, FCR, NCPR (sliding window size of 10) were obtained by CIDER^34^. CatGranule^35^, PScore^36^, ParSe^37^ and FuzDrop^38^ were used to evaluate LLPS propensity. The composition profile was obtained using Composition Profiler^39^. The charge-hydropathy plot^40^ was generated by PONDR. The secondary structure of AGARP was predicted by AlphaFold2^31^ using ColabFold^32^ and only those predictions with pLDDT >70 were analyzed.

### Statistics

Data analysis was performed by linear or nonlinear least-squares regressions using OriginPro 2023 (OriginLab). Discrimination between models was based on Akaike’s Information Criterion or Snedecor’s *F*-test. Error calculus was performed according to propagation rules based on partial derivatives of all variables.

## Data availability

All data necessary to interpret, verify and extend the research are included within the manuscript and Supplementary Information. Raw data are available on request from the corresponding author.

## Acknowledgements

We thank Prof. Joanna Trylska for the access to the CD laboratory at the Centre of New Technologies, University of Warsaw and Prof. Jarosław Stolarski for inspiring discussions. The work was supported by Polish National Science Centre grant no. 2016/22/E/NZ1/00656 to AN. The studies were performed in the NanoFun laboratories co-financed by ERDF within the Innovation Economy Operational Program POIG.02.02.00-00-025/09.

## Author contributions

B.P.K. and A.N. designed and performed the experiments, analyzed and interpreted the data and wrote the manuscript with contributions from other authors. B.P.K. and A.M cloned and expressed the proteins. T.W. and B.P.K. acquired the SEM and EDS data. B.P.K prepared the illustrations. A.N. conceived, designed and supervised the project.

## Additional information

Supplementary information is available as an additional file.

Correspondence and requests for materials should be addressed to A.N.

