## Supplementary Information for "Guiding calcium carbonate formation *via* liquid phase separation of extremely charged coral protein AGARP"

```
A.mil_AGARP          SPLRNRFNEDHDEFSKDDMA-RESFDTEEMYNAFLNRRDSSESQLEDHLLSHAKPLYDDF        59
A.mur_AGARP          SPLRNRFNEDHDEFSKDDMA-RESFDTEEMYNAFLNRRDSSESQLEDHLLSHAKPLYDDF        59
A.cer_AGARP          SPLRDRFNKDRDEFSRDDVA-RESFDTEEMYNAFLNRRDSSESQLEDHLLSHAKPLYDDF        59
A.dig_AGARP          SPLRNRFNEDHDEFSKDDMA-RESFDTEEMYNAFLNRRDSSESQLEDHLLSHAKPLYDDF        59
M.fol_AGARP-l        SPLRNRFSEDFDDFSRETMPAKESSDVVDFYDAYLRRREPSESRLEDYLLSQPKPMEDET        60
M.cap_AGARP-l        SPLRNRFSEDFDDFSRETMPAKEISDVVDFYDAYLRRREPSESRLEDYLLSQPKPMEDET        60
                     ****:**.:* *:**:: :  :*  *. ::*:*:*.**: ***:***:***: **: *:

A.mil_AGARP          FPKDTSPDDDEDSYWLESRNDDGYDLAKRKRGYDDEEAYDDFDEVDDRADDEGARDVDES        119
A.mur_AGARP          FPKDTSPNDDEDSYWLESRNDDGYDLAKRKRGYDDEEAYDDFDEVDDRADDEGARDVDES        119
A.cer_AGARP          VPKDTSPDDDEDSYWLESRNDDGYDLAQRKRGYDDEEAYDDFDEVEDRADDEGARDVDES        119
A.dig_AGARP          FPKDTSPDDDEDSYLLESRNDDGYDLAQRKRGYDDEEALDDFDEVDDRADDEGARDIDES        119
M.fol_AGARP-l        YSRE-IPDDDEDSYWGESRNDDSYDFAQRKRFYDDEDAANNKFEEANYRGYTNDADVEES        119
M.cap_AGARP-l        YSRE-IPDDDEDSYWGESRNDDSYDFAQRKRFYDDKDAANNKFEEANYRGYTNDADVEES        119
                       ::  *:******  ******.**:*:*** ***::* ::  *  :  .  .  *::**

A.mil_AGARP          DFEEDDKLPAEEESKNDMDEETFEDEPEEDKEEAREEFAEDERADE-------------        165
A.mur_AGARP          DFEEDDKLPAEEESKNDMDEETFEDEPEEDKEEAREEFAEDERADE-------------        165
A.cer_AGARP          DFDEDDKLPAEEESKNDMDEETFEDEPEEDREEAREEFAEDERADE-------------        165
A.dig_AGARP          DFDEDDKLPAEEESKNDMDEETFEDEPEEEREEAREEFAEDERADE-------------        165
M.fol_AGARP-l        L-YDEEELPDEGEGRSDMNEEAFEDDLEEDNEDANQEFNDDYEDVDPKEASDEDEQEEEE        178
M.cap_AGARP-l        L-YDEEELPDEGEGRSDMNEEAFEDDLEEDNEDANQEFNDDYEDVDPKEASDEDEQEEEE        178
                        ::::** * *.:.**:**:***: **:.*:*.:** :* .  :

A.mil_AGARP          -----REDDDADFDFN-DEEDEDEVDNKAESDIFTPEDFAGVSDEAMDNFRDDNEEEYAD        219
A.mur_AGARP          -----REDDDADFDFN-DEEDEDEVDNKAESDIFTPEDFAGVSDEAMDNFRDDNEEEYAD        219
A.cer_AGARP          -----REDDDADFDFN-DEEDEDEVDNKAESDIFTPEDFAGVSDEAMDNFRDDREEEYAD        219
A.dig_AGARP          -----REDDDADFDFN-DEEDEDEVDNKAESDIFTPEDFAGVSDEAMDNFRDDNEEEYAD        219
M.fol_AGARP-l        EEEEEEEEDEPEMDINYATEDEAKMDNNEEDDIMTSEEFENSNDEAKEKYTGDFLD----        234
M.cap_AGARP-l        EEEEEEEEDEPEMDINYATEDEAKMDNNEEDDIITSEEFENSNDEAKEKYTGDFLD----        234
                          .*:*: ::*:*   *** ::**: *.**:* *:* . .*** ::: .*  :

A.mil_AGARP          ESDDEAEEDSEETADDFEDDPEDESDETFRDEVEDESEENYQDDTEEGSEIKQNDETEEQ        279
A.mur_AGARP          ESDDEAEEDSEETADDFEDDPEDESDETFRDEVEDESEENYQDDTEEGSEIKQNDETEEQ        279
A.cer_AGARP          ESDDEAEEDSEETAEDFEDDPEDESDETFRDEVEDESEENYRDDTEDGSEIKQNDETEEQ        279
A.dig_AGARP          ESDDEAEEDSEETADDFEDDPEDESDETSRDEVEDESEENYRDDTEDGSEIRQNDETEEQ        279
M.fol_AGARP-l        E--------SEEAEEGYNDEPEEETEEEPD-KTAEDSEDESEDETEDESEEKYQDETDEE        285
M.cap_AGARP-l        E--------SEEAEEGYNDEPEEETEEEPD-KTAEDSEDESEDETEDESEEKYQDETDEE        285
                     *        ***: :.::*:**:*::*    :. ::**:: .*:**: ** : :***:*:

A.mil_AGARP          PEKKFDADKEHEDAPEPLKEKLSDESKARAEDESDKSEDAAKEIKEPEDAVEDFEDGAKV        339
A.mur_AGARP          PEKKFDADKEHEDAPEPLKEKLSDESKARAEDESDKSEDAAKEIKEPEDAVEDFEDGAKV        339
A.cer_AGARP          PEKKFDADIEHEDAPEPLKEKLSDESKARAEDESDKSEDAAKEIKEPEDAVEDFEDGAKV        339
A.dig_AGARP          PEKKFDADKEHEDAPEPLKEKLSDESKARAEDESDKSEDAAKEIKEPEDAVEDFEDGAKV        339
M.fol_AGARP-l        PEEAFDEIEYPEDDPEPLKAKLSDEIQRIRYQ--------NDDLKDSEDEVQDFDESEMV        337
M.cap_AGARP-l        PEEAFDEIEYPEDDPEPLKAKLSDEIQGKRYQ--------NDDLKDSEDEVQDFDESEMV        337
                     **: **     ** ***** ***** :    :         .::*: ** *:**::.  *

A.mil_AGARP          SEDEAELLDDEAELSDDEAELSKDEAEQSSDEAEKSEDKAEKSE--DEAELSEDEAKQSE        397
A.mur_AGARP          SEDEAELLDDEAELSDDEAELSKDEAEQSSDEAEKSEDKAEKSE--DEAELSEDEAKQSE        397
A.cer_AGARP          SEDEAELLDDEAELSDDEAELSKDEAEQSSDEAEKSEDKAEKSE--DESELSEDEAKQSE        397
A.dig_AGARP          SEDEAELLDDEAELSDDEAELSKDEAEQSSDEAEKSEDKAEKSEFEDEAELSEDEAKQSE        399
M.fol_AGARP-l        NKEDITR----------------EEA----------------------------------        347
M.cap_AGARP-l        NKEDITR----------------EEA----------------------------------        347
                     .:::                   :**

A.mil_AGARP          DEAEKAEDAAGKESNDEGKKREDEAVKSKGIARDESEFAKAKKSNLALKRDENRPLAKGL        457
A.mur_AGARP          DEAEKAEDAAGKESNDEGKKREDKAVKSKGIARDESEFAKAKKSNLALKRDENRPLAKGL        457
A.cer_AGARP          DEAEKAEDAAGKVSNDKGKKREDEAVKSKGIARDESEFAKAKKGNLALKRDENRLLAKGL        457
A.dig_AGARP          DEAEKAEDAAGKESNDEGKKREEEAVKSKGIARDESEFAKAKKSNLALKRDENRLLAKGL        459
M.fol_AGARP-l        -----------------------------------------KSDASLKQRDERYQPAKRL        366
M.cap_AGARP-l        -----------------------------------------KSDASLKQRDERYQPAKRL        366
                                                              *..    :***.   ** *

A.mil_AGARP          RESAAHLRDFPSEKKSKDAAQGNIENELDYFKRNAFADSKDA-EPYEFDK        506
A.mur_AGARP          RESAAHLRDFPSEKKSKDAAQGNIENELDYFKRNAFADSKDA-EPYEFDK        506
A.cer_AGARP          RESAAHLRDFPSDKKSKDAAQGNIENELDYFKRNAFADSKDA-EPYEFDK        506
A.dig_AGARP          RESAAHLRDFPSEKKSKDAAQGNIGNANYDEV--LIP-------------        494
M.fol_AGARP-l        HESLARLRNLPSEKNLKND-REKIEKELDYFKKNTLPNKANVEDSYVFAK        415
M.cap_AGARP-l        HESLARLRNLPSEKNLKND-REKIEKELDYFKKNTLPNKANVEDSYVFAK        415
                     :** *:**::**:*: *:  : :* :         :
```

**Fig. S1**. Multiple sequence alignments of AGARP from *Acropora millepora* (A.mil), *A. muricata* (A.mur), *A. cervicornis* (A.cer), putative AGARP from *A. digitifera* (A.dig), AGARP-like (AGARP-l) from *Montipora foliosa* (M.fol) and from *M. capricornis* (M.cap), which were identified as similar through a BLASTP search. Sequence identity to *A. millepora* AGARP is 99.60, 95.85, 95.33, 45.04 and 44.53%, respectively. All sequences are given without signal peptide. Code: * , identical amino acid (aa) residue; : , aa of strongly similar properties; . , aa of weakly similar properties.

Both AGARP and CARP1-3[1] contain multiple short negatively charged blocks formed by adjacent Asp and Glu residues that may serve as calcium-chelating motifs. However, the longest such block in AGARP contains seven acidic residues, in contrast to CARP3, which has up to twenty four D/E repeats in a continuous sequence.

```
AGARP       SPLRNRFNEDHDEFSKDDMARESFDTEEMYNAFLNRRDSSESQLEDHLLSHAKPLYDDFF      60
CARP1       ------------------------------------------------------------      0

AGARP       PKDTSPDDDEDSYWLESRNDDGYDLAKRKRGYDDEEAYDDFDEVDDRADDEGARDVDESD      120
CARP1       ------------------------------------------------------------      0

AGARP       FEEDDKLPAEEESKN-------DMDEETFEDEPEEDKEEAREEFAEDERADEREDDDADF      173
CARP1       -EGDHLKPGHSEDEHDEDEHDEEMADHADEQNPADEEETEDEEKDDDKMEDDSDDDEEDE      59
             * *.  *...*.::       :* :.: *::* :::*   **  :*:  *: :**: *

AGARP       DFNDEEDEDEV-DNKAESDIFTPED---FAGVSDEAMDNFRDDNEEEYADESDDEAEEDS      229
CARP1       SQGDDEGEDENDQSHLEHDAFLGDNYTEFKGLSPEDAKT-------KLAQLIKDEVDLNK      112
            . .*:*.***  :.: * * *  ::   * *:* *  ..       : *:  .**.: :.

AGARP       EETADDFEDDPEDESDETFRDEVEDESEENYQDDTEEGSEIKQNDETE-------EQPEK      282
CARP1       ----DGV--LTEDEIRQRFHVT-----TKEYRKKE-VMETMKQHDEDKDGKVSWEEFKKG      160
                *..    ***  : *:        ::*:..    . :**:** :       *  :

AGARP       KFDADKEHEDAPEPLKEKLSDESKARAEDESDKSEDAAKEIKEPEDAVEDFEDGAKVSED      342
CARP1       HFSDDGKDEDAKEQMKEDE---EKFKFADED---GDGKLDL---EEYMAFYHPGDNPRMT      211
            :*. * :.*** * :**.    .* :  **.    *.  ::   *: :  :. * :

AGARP       EAELLDDEAELSDDEAELSKDEAEQSSDEAEKSEDKAEKSEDEAELSEDEAKQSEDEAEK      402
CARP1       EFTI-------ED---SLKKH-------------D-------------------------      223
            *  :       .*   .*.*.             *

AGARP       AEDAAGKESNDEGKKREDEAVKSKGIARDESEFAKAKKSNLALKRDENRPLAKGLRESAA      462
CARP1       --------KDKDGQVSKKEFLATFSDVNDDAKEEMEKDFNNNFDKDKNGRLNKE----EM      271
                    .:.:*:  :.* : : . ..*:::    *. *  :.:*:*  * *

AGARP       HLRDFPSEKKSKDA-----------AQGN-----IENELDYFKRNAFADSKDAEPYEFDK      506
CARP1       KSWLFPDDDFSTEEPKTLIKEADEDKDGKLTMDEIMKNYKVFIEDEPEDSSHD---EL--      326
            :   **.:. *.:             :*:     * :: . * .:   **..    *:
```

```
AGARP       SPLRNRFNEDHDEFSKDDMARESFDTEEMYNAFLNRRDSSESQLEDHLLSHAKPLYDDFF      60
CARP2       APVEN----------------------------------------------------EI-      7
            :*:.*                                                    ::

AGARP       PKDTSPDDDEDSYWLESRNDDGYDLAKRKRGYDDEEAYDDFDEVDDRADDEGARDVDESD      120
CARP2       ------------------------------------------------RIRGPKLED---      16
                                                          .* :  *

AGARP       FEEDDKLPAEEESKNDMDEETFEDEPEEDKEEAREEFAEDERADEREDDDADFDFNDEED      180
CARP2       -EEEGNFPPIMPAQFELKEREFPKKEEERKE-----AKEDENM----LREELKHFRDEES      66
             **:.::*    :: ::.*. * .: ** **       ***.       :   .*.***.

AGARP       EDEVDNKAESDIFTPEDFAGVSDEAMDNFRDDNEEEYADESDDEAEEDSEETADDFEDDP      240
CARP2       LKNVITRLEREL----AFEKTE-------REENR-ETEDLSNEELV--ERELPEEVDEIP      112
             .:* .: * ::     *  ..       *::*. *  * *::*    ..*  ::.:: *

AGARP       EDESDETFRDEVEDESEENYQDDTEEGSEIKQNDETEEQPEKKFDADK----EHEDAPEP      296
CARP2       EEKGARELKEEN--GLEMFYRN-LQR--KLKEK-QERDMPVKEMEHESPEDQEEEMQERE      166
            *::. . :::*     *  *::  :.  ::*:: : .: * *::: :.    *.*   .

AGARP       LKEKLSDESKARAEDESDKSEDAAKEIKEPEDAVEDF-EDGAKVSEDEAELLD---DEAE      352
CARP2       LDEEFKEKSKRELEE-------------------EDLEETGAEEREDKRELAEEVSSREE      207
            *.*::.::** . *:                   **: * **:  **: ** :   .. *

AGARP       LSDDEAELSKDEAEQSSDEAEKSEDKAEKSEDEAELSEDEAKQSEDEAEKAEDAAGKESN      412
CARP2       LEENEEELALKRKRG--------E---ENMATEWEIPESVEH--YDENKRSKH-PPKHMR      253
            *.::* **: .. .         *   *:   * *: *.  :   **  ::::.   *. .

AGARP       DEGKKREDEAVKSKGIAR-DESEFAKAKKSNLALKRDENRPLAKGLRESAAHLRDFPSEK      471
CARP2       EREAERERERFDDHGHKEREREEFR-ERQRELALSN--------GG---KLHERELEGRK      301
            :.  :** * ...:*  . :..**   :: :***..        *      * *:: ..*

AGARP       KSKDA-AQGNIENELDYFKRNAFADSKDAEPYEFDK---      506
CARP2       QRQEIGLHGVRREE--------------SERFRFRVRGE      326
            : ::   :*  .:*              :* :.*
```

```
AGARP      SPLRNRFNEDHDEFSKDDMARESFDTEEMYNAFLNRRDSSESQLEDHLLSHAKPLYDDFF      60
CARP3      ------------------MPAEH------YGSYSQN---------IMLLDTHEDKARNYV      27
                             *  *       *.:: :.           **.  :    ::.

AGARP      PKDTSPDDDEDSYWLESRNDDGYDLAKRKRGYDDEEAYDDFDEVDDRADDEGARDVDESD      120
CARP3      PE--------------SANATDPAVAEPSEAEND-PAQSETEPAAEEASTDS-----AAD      67
           *:              * *  .  :*: ... :*  * .: : . :.*. :.      :*

AGARP      FEEDDKLPAEEESKNDMDEETFEDEPEEDKEEAREEFAEDERADEREDDDADFDFNDEED      180
CARP3      TKEDDSAAADDSSDDDLDDDSVDEN-------------------------------DEDD      96
            :***.  *::.*.:*:*:::.:::                               **:*

AGARP      EDEVDNKAESDIFTPEDFAGVSDEAMDNFRDDNEEEYADESDDEAEEDSEETADDFEDDP      240
CARP3      EDDEDDE----------------------------DDEDDEDDEDDEDDEDDGDDSD-DG      127
           **: *::                            :  *:.*** :**.*: .** : *

AGARP      EDESDETFRDEVEDESEENYQDDTEEGSEIKQNDETEEQPEKKFDADKEHEDAPEPLKEK      300
CARP3      DDESD---------GDDENDGDDEDDGDDE------------------------------      148
           :****          .:**  ** ::*.:

AGARP      LSDESKARAEDESDKSEDAAKEIKEPEDAVEDFEDGAKVSEDEAELLDDEAELSDDEAEL      360
CARP3      ------------------------------------------------------------      148

AGARP      SKDEAEQSSDEAEKSEDKAEKSEDEAELSEDEAKQSEDEAEKAEDAAGKESNDEGKKRED      420
CARP3      ------------------------------------------------------------      148

AGARP      EAVKSKGIARDESEFAKAKKSNLALKRDENRPLAKGLRESAAHLRDFPSEKKSKDAAQGN      480
CARP3      ------------------------------------------------------------      148

AGARP      IENELDYFKRNAFADSKDAEPYEFDK      506
CARP3      --------------------------      148
```

---

```
AGARP      SPLRNRFNEDHDEFSKDDMARESFDTEEMYNAFLNRRDSSESQLEDHLLSHAKPLYDDFF      60
CARP4      ------------------------------------------------------------      0

AGARP      PKDTSPDDDEDSYWLESRNDDGYDLAKRKRGYDDEEAYDDFDEVDDRADDEGARDVDESD      120
CARP4      ------------------------------------------------------------      0

AGARP      FEEDDKLPAEEESKNDMDEETFEDEPEEDKEEAREEFAEDERADEREDDDADFDFNDEED      180
CARP4      -----------------------------------MIVRDNDDDD-DDDDDDDDDDDDD-      23
                                               :..*:  *: :*** * * :*::

AGARP      EDEVDNKAESDIFTPEDFAGVSDEAMDNFRDDNEEEYADESDDEAEEDSEETADDFEDDP      240
CARP4      -----------------------------------------DDDGVDDDDDDDDDFSDDN      42
                                                    **:. :*.::  ***.**

AGARP      EDESDETFRDEVEDESEENYQDDTEEGSEIKQN--DETEEQPEKKFD-----ADKEHEDA      293
CARP4      E----EMLSFEVDELEE-----KDVDGNDVDTMQRHSVDSFDDVDFTFTKVDTQAKYDGL      93
           *    * :  **:: .*     .  :*.::.    ...:.  : .*      :: :::.

AGARP      PEP---LKEKLSD-------------ESKARAEDESDKSEDAAKEIKEPEDAVEDFEDGA      337
CARP4      PVTNVNLSATLPSSSSLEIMVYLFRRAGKVTFGNETFRVEKGTIKF-NIRISNWDFCDGS      152
           *     *. .* .              .*.   :*: : *..: :: : . :  ** **:

AGARP      KVSEDEAE---LLDDEAELS--DDEAELSKDEAEQSSDEAEKSEDKAEKSEDEAELSEDE      392
CARP4      RFDCDEGKIGEFLDLKLKIKSKDSPAEISDDDR--------KAAA-------KRPICEDE      197
           :.. **.:   :** : ::.  *. **:*.*:         *:         :  :.***

AGARP      AKQSE--D---EAEKAEDAAGKESNDEGKKREDEAVKSKGIARDESEFAKAKKSNLALKR      447
CARP4      DSDDDGVDDPTDSDNDDDDC-PSIYDIG--GDSEMLLNKGVVTDDDKYTALPEGYPKFE-      253
            .:.:  *   :::: :* .  .  * *   :.* : .**:. *:.:::   :.   ::

AGARP      DENRPLAKGLRESAAHLRDFPSEKKSKD----AAQGNI---------ENELDYFK-RNAF      493
CARP4      -----VEDGEKKFVF---RIPKFKKSVLVDPSVNVGRVNGVNSLAHANTGLAFLLFIAAF      305
                : .* :: .     :*. ***      .  *.:         :. * ::    **

AGARP      ADSKDAEPYEFDK      506
CARP4      I----VTNYV---      311
```

**Fig. S2** Alignments of the AGARP sequence with CARPs from *Stylophora pistillata*: CARP1 (Uniprot ID: R4IW69), CARP2 (R4IW51), CARP3 (R4IW63), CARP4 (M1SN56). * and cyan, identical aa residue; : , aa of strongly similar properties; . , aa of weakly similar properties; green, substitution of D by E or vice versa; grey, W/F/Y substitution.

**Table 1.** Partition coefficients, $K_{AV}$, of standard folded proteins, His$_6$-SUMO-AGARP and AGARP determined *via* SEC at 10 °C, along with Stokes radii, $R_H$, of folded proteins (from literature) and those of studied proteins (calculated *via* SEC calibration on folded proteins). Data shown in **Fig. 2b**.

| **Protein** | **$K_{AV}$** | **$K_{AV}$ err** | **$R_H$ (Å)** | **$R_H$ err (Å)** |
|---|---|---|---|---|
| Lysozyme | 0.689 | 0.007 | 19[2] | |
| α-chymotrypsinogen A | 0.5269 | 0.0002 | 22.4[3] | |
| BSA monomer | 0.354 | 0.005 | 35.5[3] | |
| BSA dimer | 0.234 | 0.004 | 43[3] | |
| HSA monomer | 0.3732 | 0.0006 | 35.1[4] | |
| Apoferritin monomer | 0.146 | 0.003 | 60.25[5] | |
| Apoferritin dimer | 0.050 | 0.003 | 88.75[5] | |
| His$_6$-SUMO-AGARP | 0.0942 | 0.0014 | 74 | 4 |
| AGARP | 0.1235 | 0.0004 | 66 | 4 |

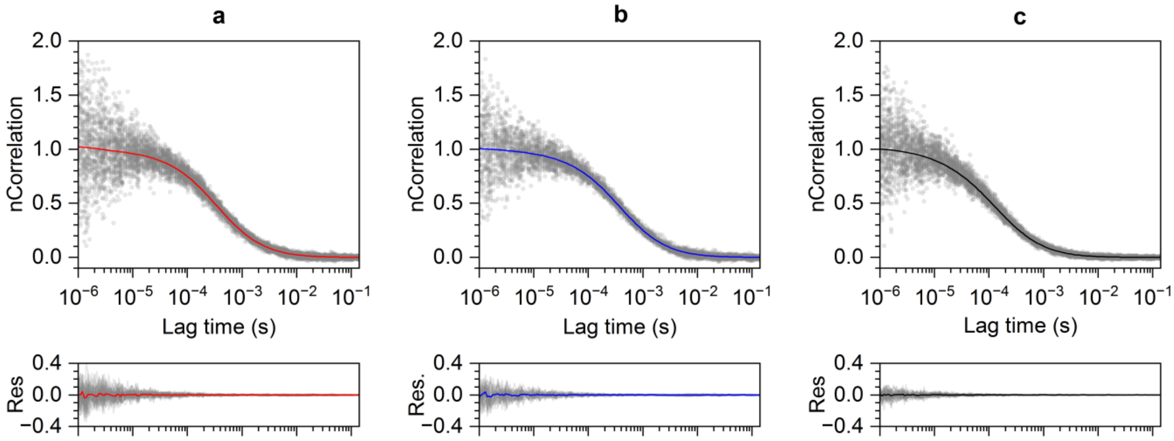

**Fig. S3** Experimental FCS points (dots) used for global fitting of the diffusion models (lines) for (**a**) AGARP, (**b**) $His_6$-SUMO-AGARP and (**c**) HSA.

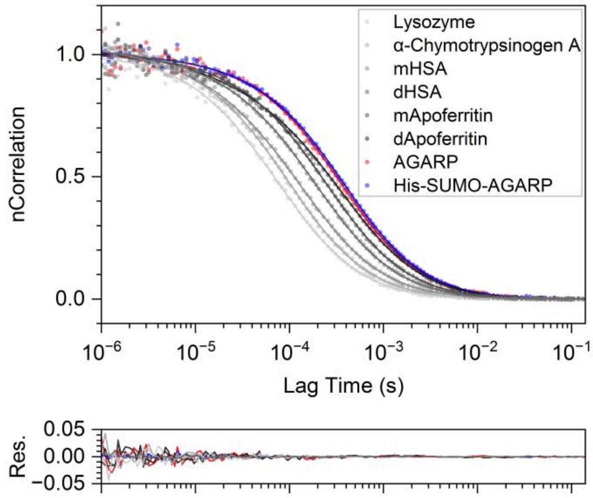

**Fig. S4** Example normalized FCS autocorrelation curves for folded proteins, AGARP and $His_6$-SUMO-AGARP (upper panel) with corresponding fit residuals (bottom panel). Dots, experimental points; lines, fitted models of diffusion.

**Table 2.** Cube root of molar mass, $M^{1/3}$ and number of amino acid residues in the protein chain, *N*, of AF488, folded proteins, $His_6$-SUMO-AGARP and AGARP together with their Stokes radii, $R_H$, determined by FCS and SEC. Data shown in **Fig. 2c**.

| | | | **FCS** | | **SEC** | |
|---|---|---|---|---|---|---|
| | **$M^{1/3}$ ($Da^{1/3}$)** | **N** | **$R_H$ (Å)** | **$R_H$ err (Å)** | **$R_H$ (Å)** | **$R_H$ err (Å)** |
| AF488 | 9.02 | | 5.4 | 0.2 | | |
| Lysozyme | 24.28 | 129 | 19.9 | 1.5 | | |
| α-chymotrypsinogen A | 29.50 | 245 | 23 | 2 | | |
| HSA monomer | 40.51 | 585 | 32 | 5 | | |
| HSA dimer | 51.04 | | 44 | 6 | | |
| Apoferritin monomer | 78.25 | 4176 | 58 | 4 | | |
| Apoferritin dimer | 98.58 | | 79 | 8 | | |
| $His_6$-SUMO-AGARP | 41.55 | 624 | 81 | 5 | 74 | 4 |
| AGARP | 38.78 | 506 | 77 | 5 | 66 | 4 |

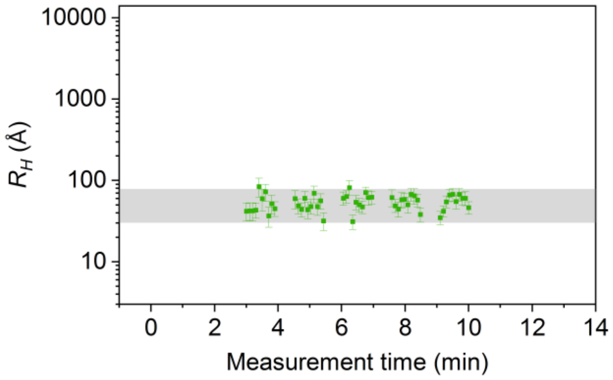

**Fig. S5** Changes of AGARP $R_H$ during FCS measurement in buffer F prior to addition of $Ca^{2+}$ ions. Shaded grey area marks 95% confidence interval for all values.

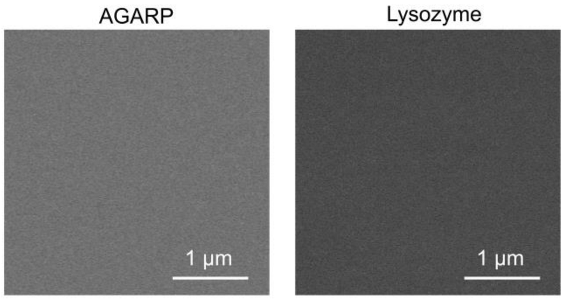

**Fig. S6** SEM images of microscopic slide after incubating on it AGARP and lysozyme for 10 min (prior to addition of calcium ions).

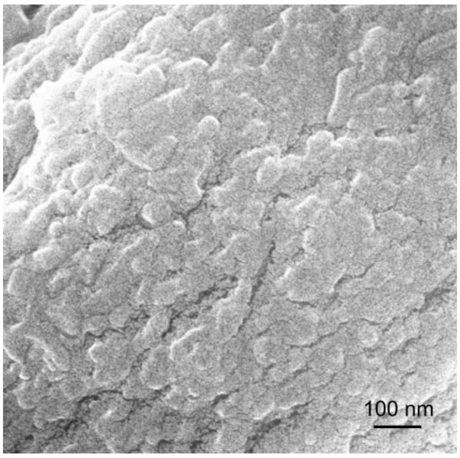

**Fig. S7** SEM image of the fragment of the phase formed after the FCS experiment with 200 nM AGARP and 5 mM $CaCl_2$ analyzed by EDS at 10 kV accelerating voltage.

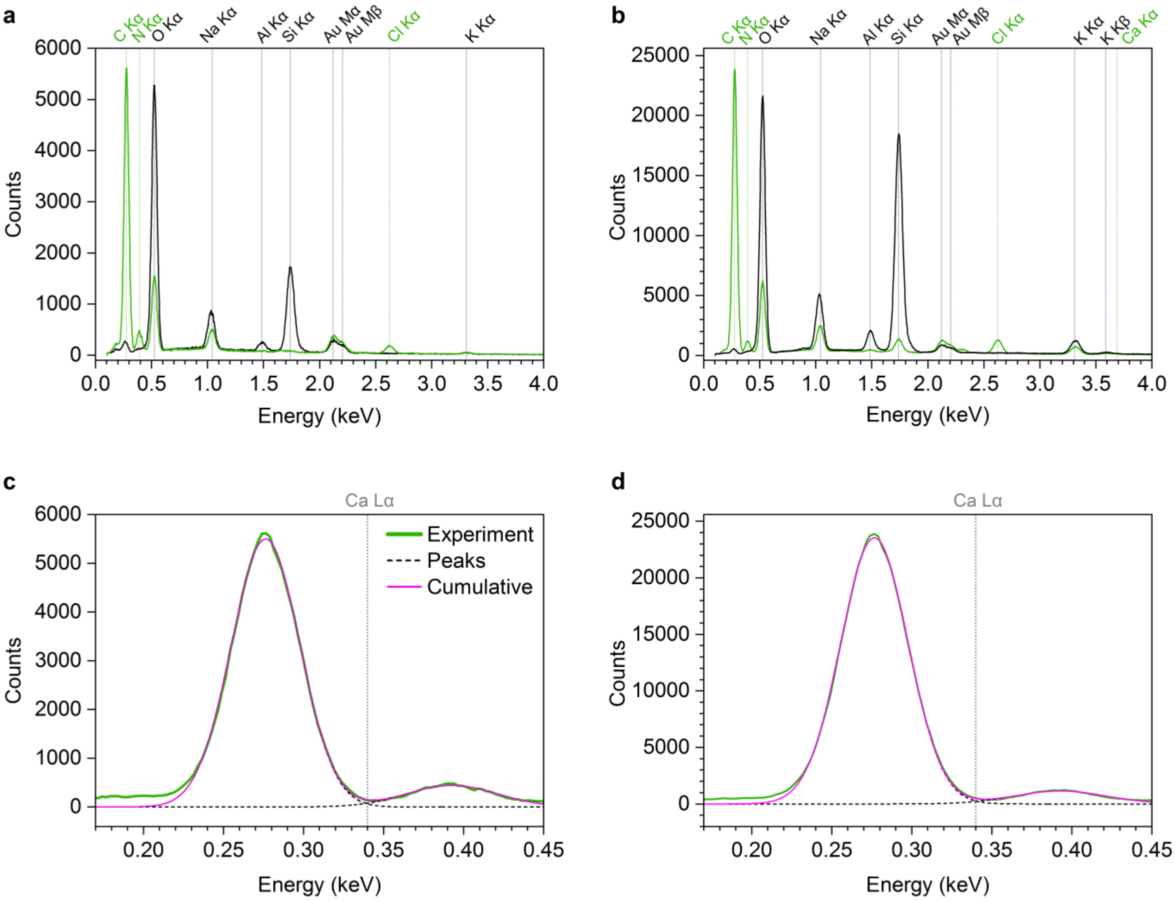

**Fig. S8** EDS spectra of $CaCO_3$ phase containing AGARP (green) and glass slide as a control (black) obtained at 5 kV (**a, c**) and 10 kV (**b, d**) accelerating voltage; (**a, b**) observed lines of elements marked as green and black vertical dotted lines; (**c, d**) Gaussian functions were fitted to the experimental peaks of C Kα and N Kα (black broken lines) and the cumulative spectra are shown (magenta); gray vertical dotted line marks the position of the absent Ca Lα line at 0.34 keV.

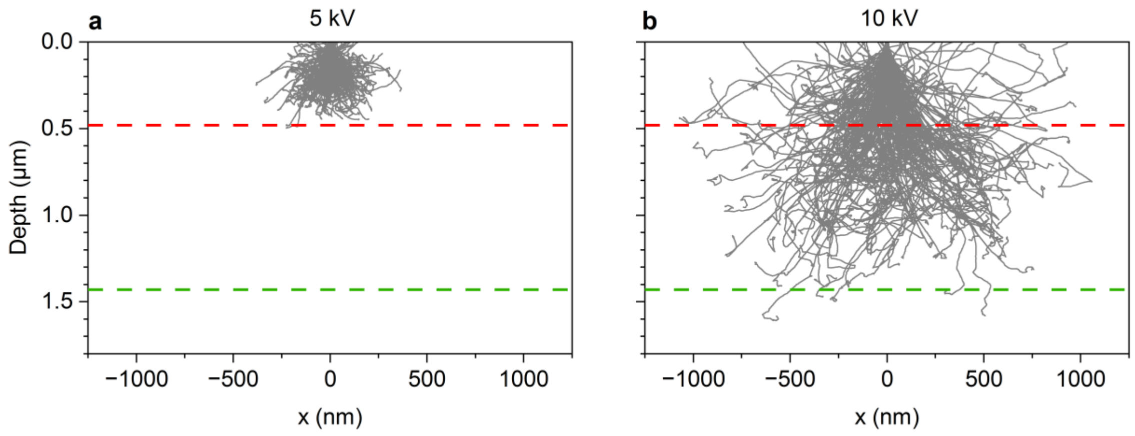

**Fig. S9** Monte Carlo simulations of electron trajectories simulated in CASINO v2.4.8.1[6] for 10 nm beam radius at (**a**) 5 kV and (**b**) 10 kV accelerating voltage. The maximum limits of the thickness of a $Ca^{2+}$-containing layer are indicated by red and green dashed lines. Default settings were applied; multilayer sample was analyzed: layer 1 (480 nm) contained C, N, O, Na, Cl, layer 2 (1430 nm): contained additionally Ca, substrate: Al, O, Si and K.

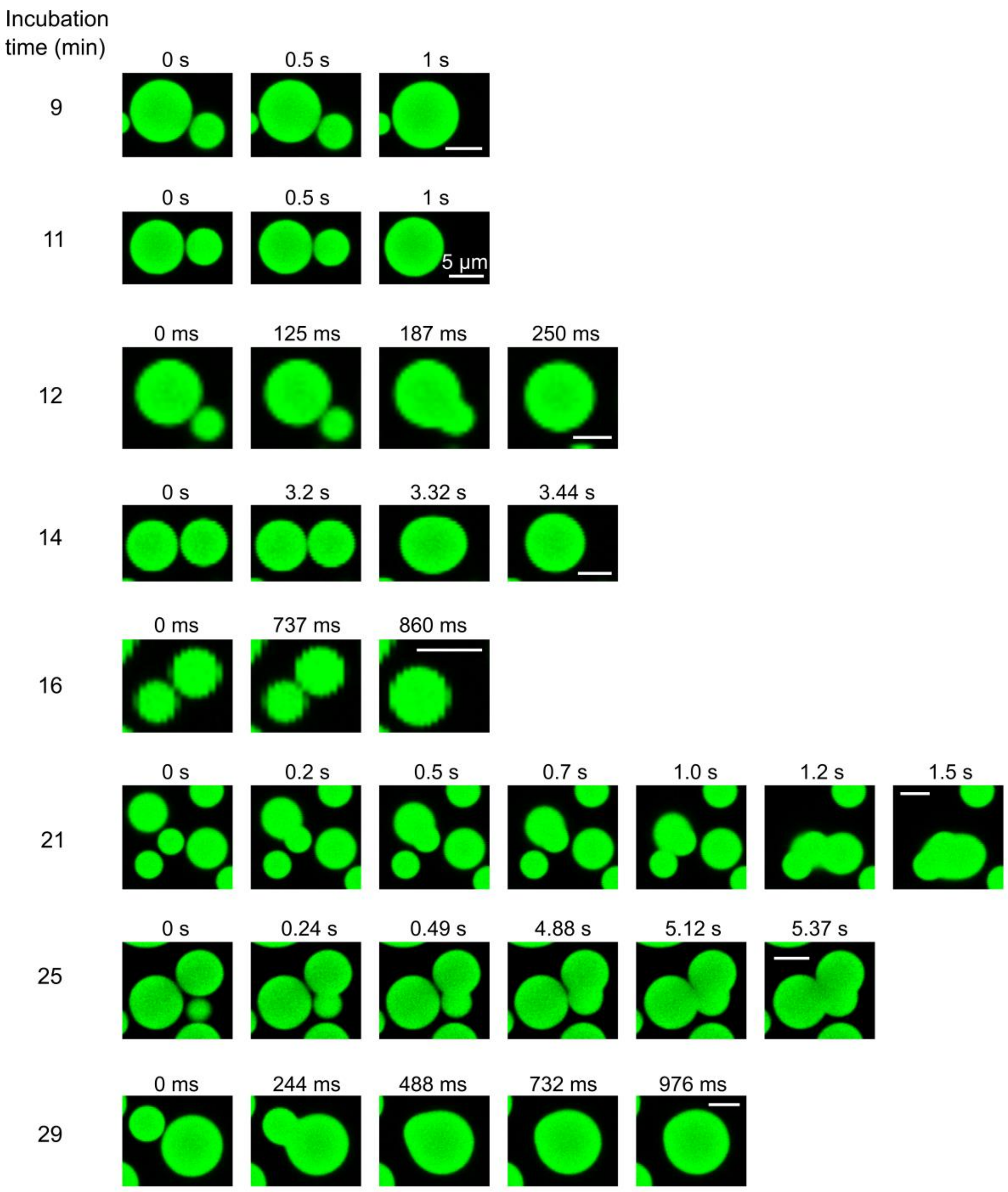

**Fig. S10** Fluorescence confocal microscopy images showing coalescence of droplets with respect to incubation time at 50 μM AGARP, 100 mM $CaCl_2$ and 7% PEG 4k on the surface of the slide passivated with Silanization solution I, Selectophore™ (Merck) and then Pluronic F-127 (Molecular Probes). Bars are 5 μm in each panel.

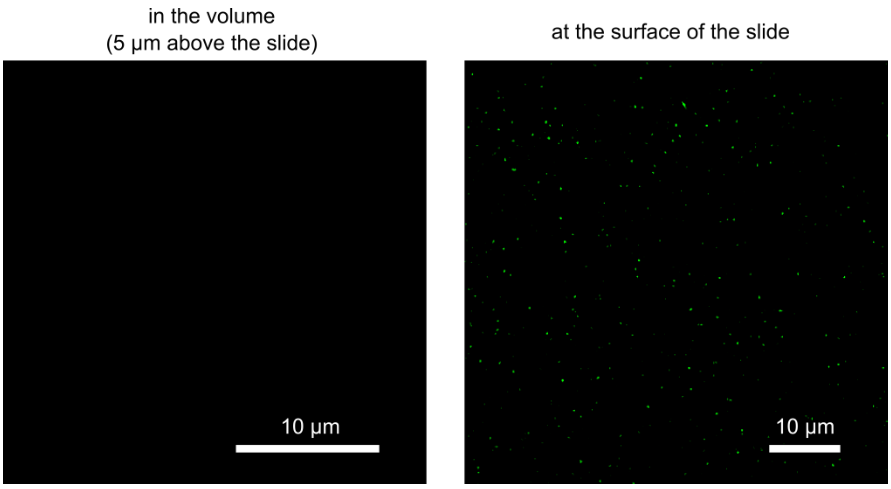

**Fig. S11** Fluorescence confocal microscopy images showing no droplets in the volume and aggregates on the surface of the slide formed at 50 µM AGARP, 100 mM $CaCl_2$ and no PEG 4k.

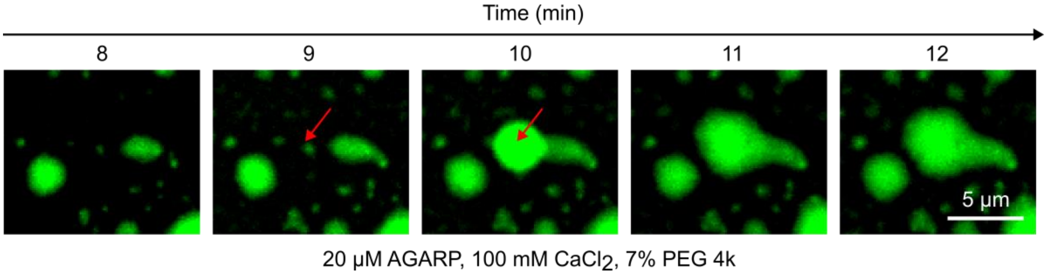

**Fig. S12** Fluorescence confocal microscopy images showing the droplets sinking by gravity and accumulation of the dense phase on the surface of the slide.

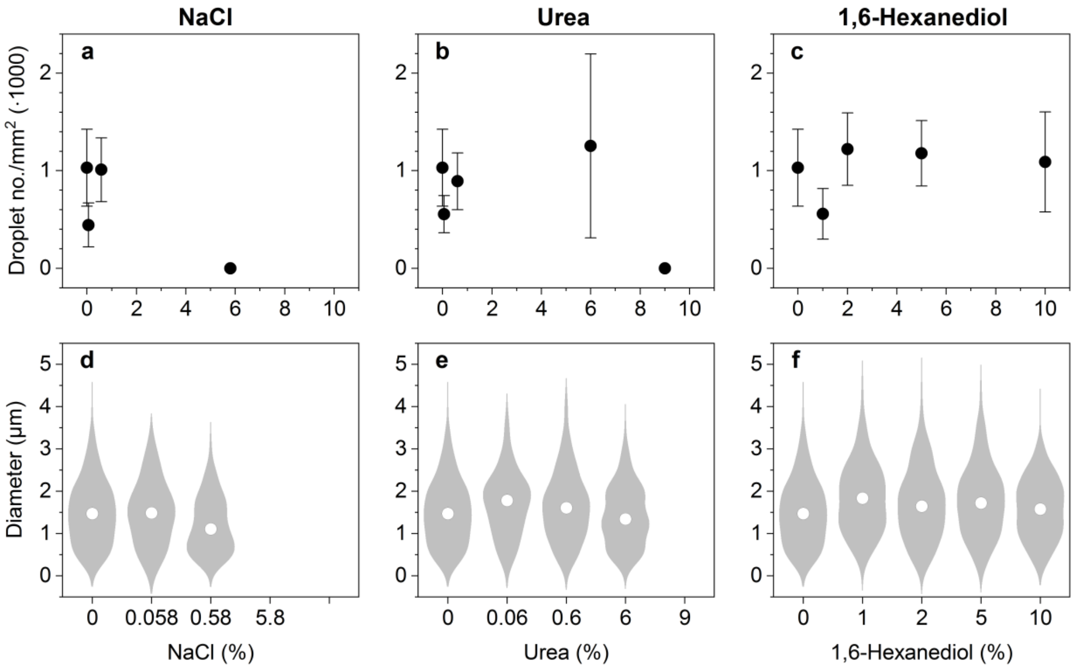

**Fig. S13** Changes of number (**a-c**) and diameter (**d-f**) of droplets formed in the volume (5 µm above the slide) at 5 µM AGARP, 50 mM $CaCl_2$, 10% PEG 4k with increasing concentration of NaCl, urea and 1,6-hexanediol.

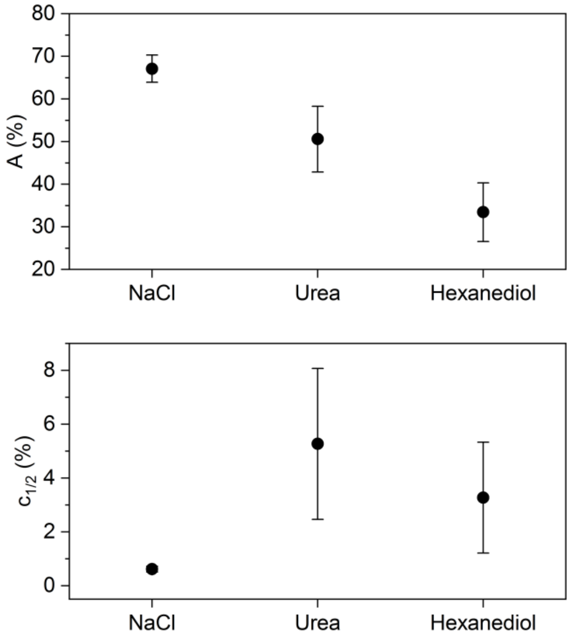

**Fig. S14** Parameters in the eq. 5 fitted to the data showing the dependence of the percentage area occupied by the liquid phase on the concentration of NaCl, urea and 1,6-hexanediol.

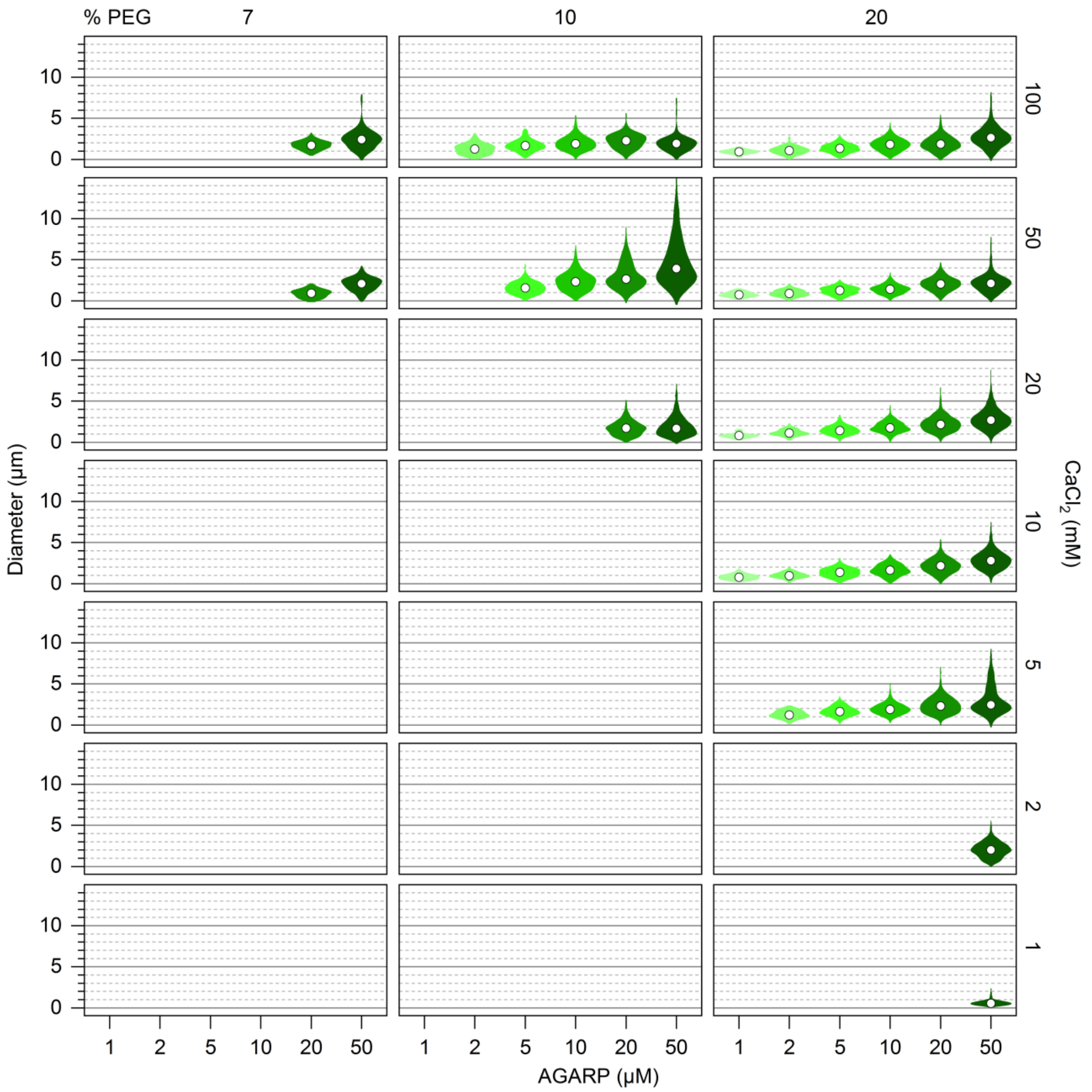

**Fig. S15** Diameter distribution of AGARP-containing droplets. Increasing intensity of green color indicates increasing concentration of AGARP.

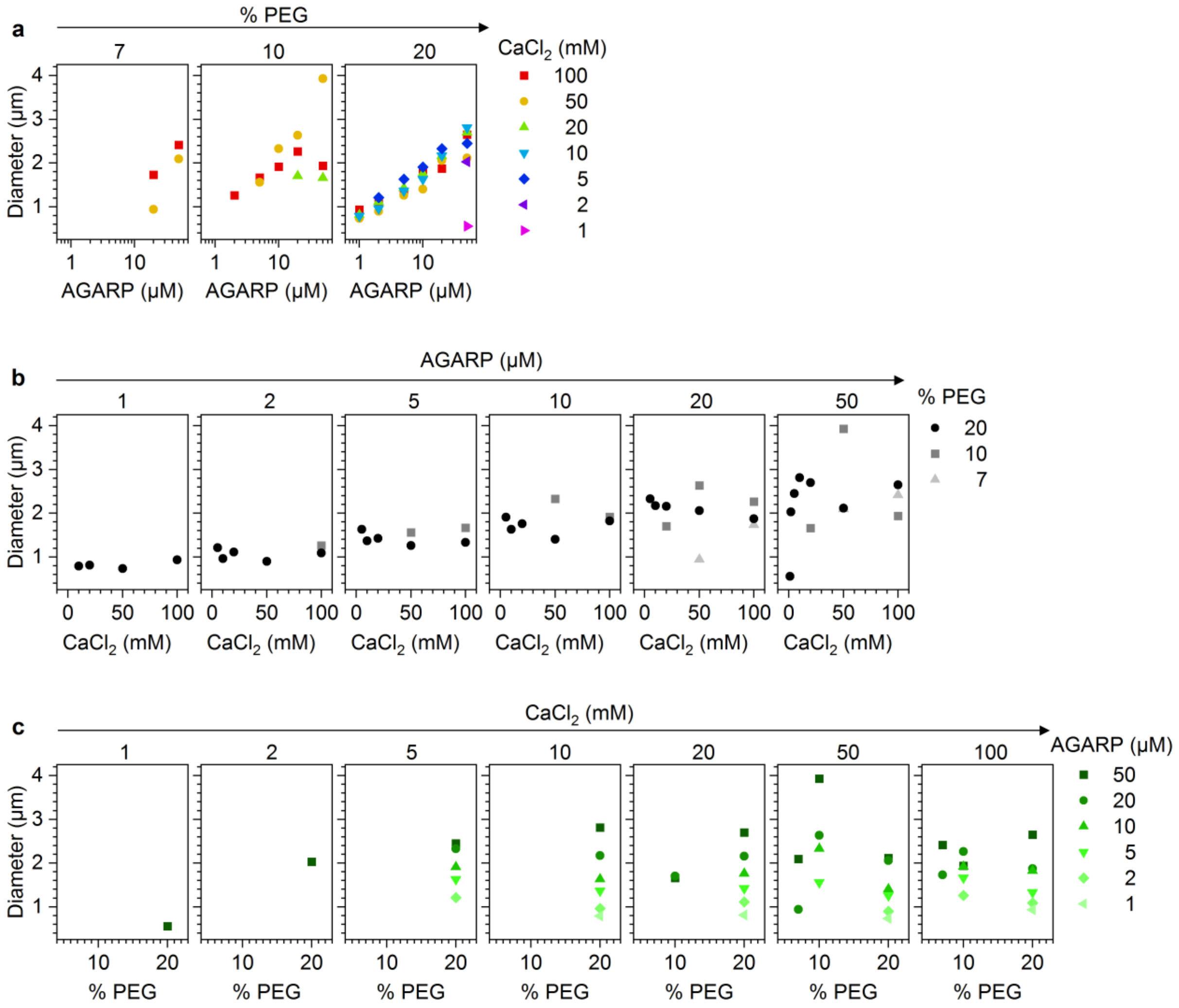

**Fig. S16** Median diameter of droplets as a function of (**a**) AGARP, (**b**) $CaCl_2$ and (**c**) PEG concentration.

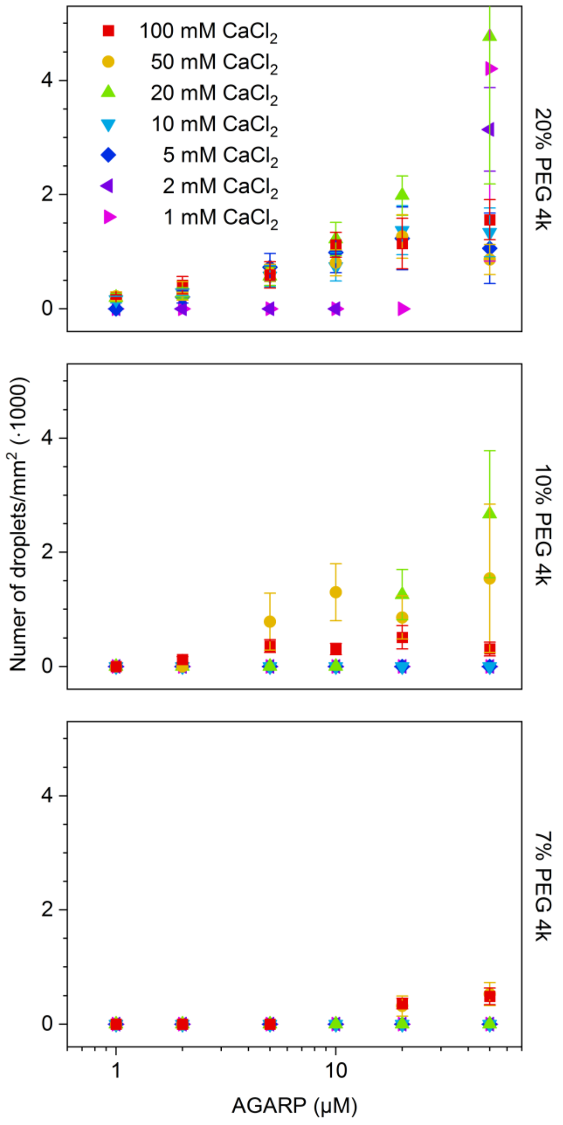

**Fig. S17** Plots showing number of droplets in the volume (5 um above the slide) per $mm^2$ as a function of AGARP concentration at varying $CaCl_2$ and PEG 4k concentrations.

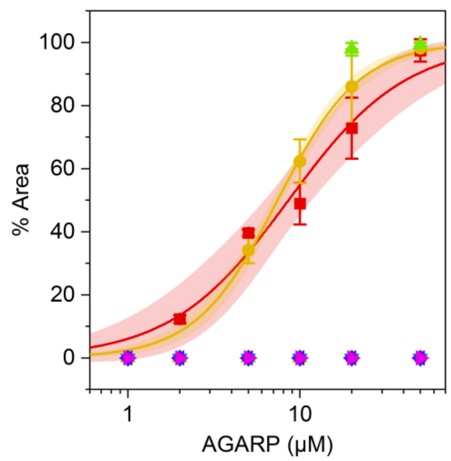

**Fig. S18** Percentage of area occupied by AGARP-rich phase at 10% PEG 4k as a function of AGARP concentration. $CaCl_2$ (mM): 100 (red squares), 50 (orange circles), 20 (green up-pointing triangles), 10 (blue down-pointing triangles), 5 (navy blue diamonds), 2 (purple left-pointing triangles), 1 (magenta right-pointing triangles), 0 (black hollow circles). **Eq. 4** was fitted to experimental data (lines colored correspondingly). Shaded area, 95% CI.

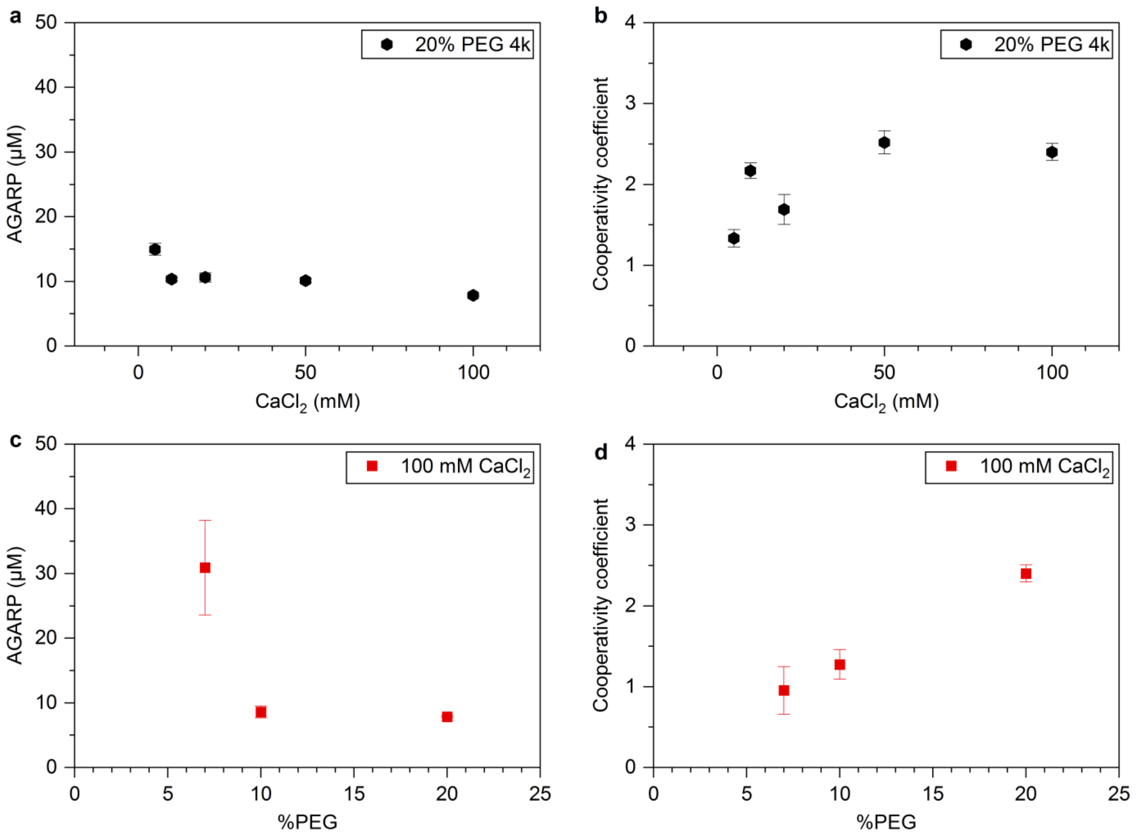

**Fig. S19** Plots showing fitted parameters: *k* [AGARP (µM)] and *n* [cooperativity coefficient] in eq. 4 for maximal PEG and $CaCl_2$ concentrations as a function of (**a**, **b**) PEG 4k and (**c**, **d**) $CaCl_2$ concentration.

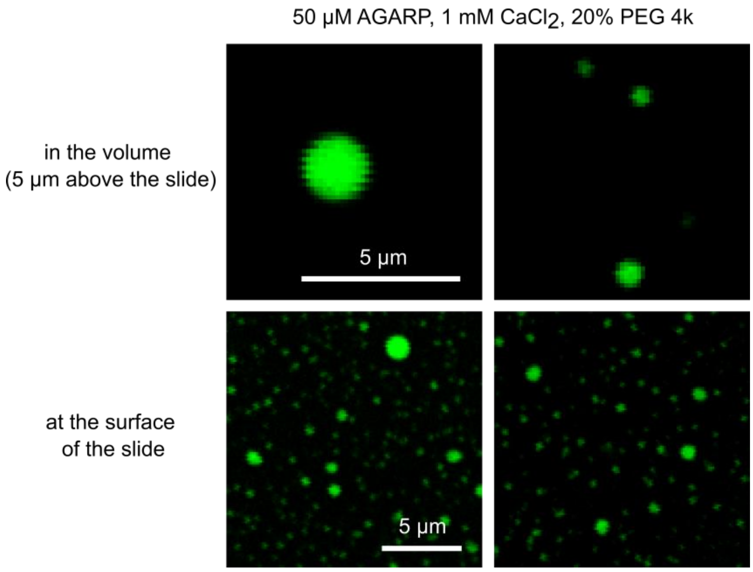

**Fig. S20** Magnification of AGARP-containing droplets and liquid phase shown in **Fig. 5a** and **e** formed at 50 µM AGARP, 1 mM $CaCl_2$ and 20% PEG 4k.

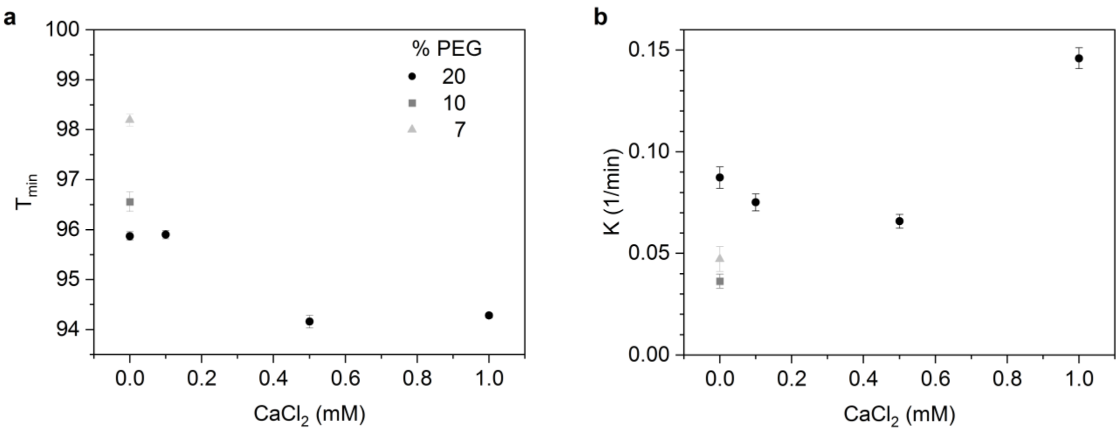

**Fig. S21** Parameters in the eq. 6 fitted to the data showing the dependence of the transmission difference at 330 nm on time in **Fig. 5d**.

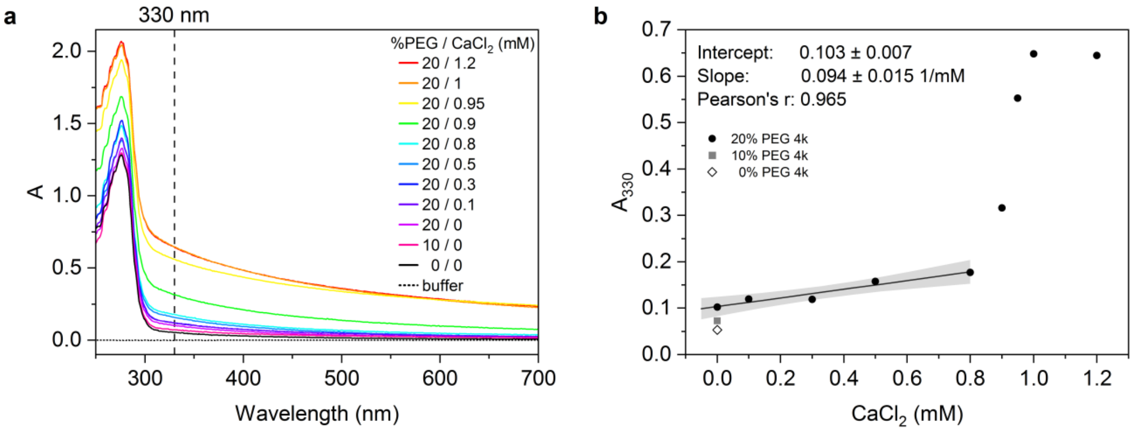

**Fig. S22 a** Absorption spectrum of 50 µM AGARP with different concentrations of PEG 4k and $CaCl_2$ measured 10 min after mixing; **b** Dependence of absorbance at 330 nm (marked with black dashed line in **a**) on $CaCl_2$ concentration. Linear function was fitted to 0-0.8 mM range.

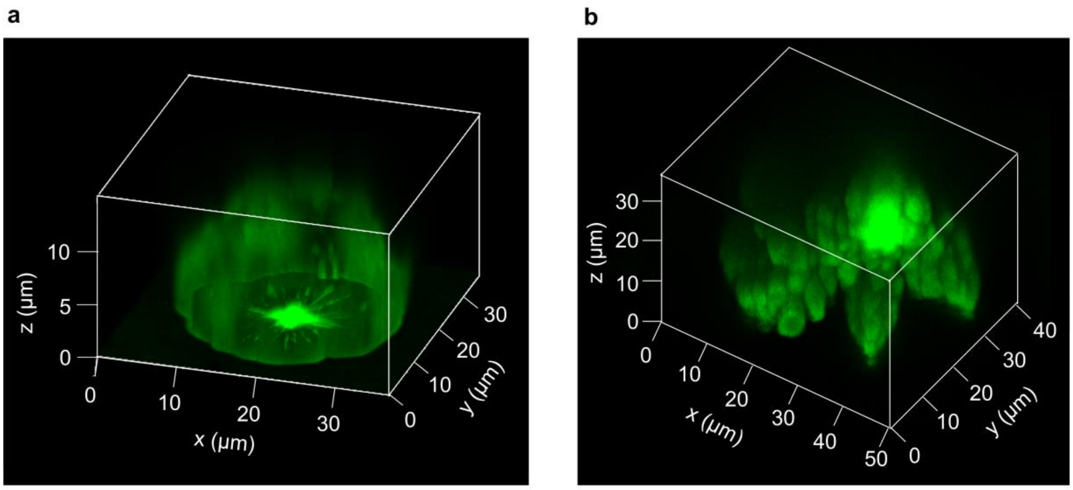

**Fig. S23** Z-stacks of $CaCO_3$ phases formed with 100 mM $CaCl_2$, 1 µM AF488-labelled AGARP and 20% PEG 4k in buffer A with (**a**) 0.4 mg (12.5 mg/$cm^3$) and (**b**) 2 mg (62.5 mg/$cm^3$) of solid $(NH_4)_2CO_3$ as a source of $CO_2$.

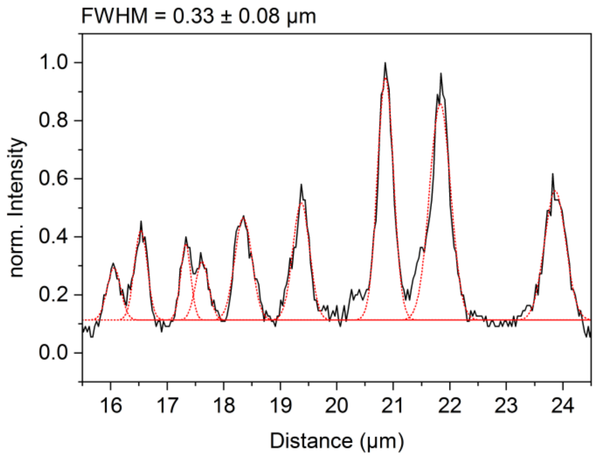

**Fig. S24** Profile of fibers visible within $CaCO_3$ phases formed with 100 mM $CaCl_2$, 1 µM AF488-labelled AGARP and 20% PEG 4k in buffer A with 0.4 mg (12.5 mg/$cm^3$) of solid $(NH_4)_2CO_3$ as a source of $CO_2$ shown in **Fig. 8a**. Black line, fibers profile; red line, fitted Gaussian peaks; FWHM, full width at half maximum.

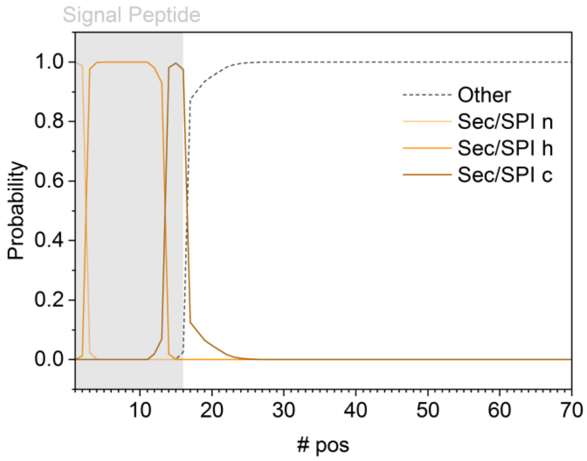

**Fig. S25** Prediction of the presence and location of the signal peptide (SP) in the AGARP sequence using SignalP 6.0[7]. SP is marked in grey. The acronym Sec/SPI indicates that the predicted SP is Sec translocon[8] (located in endoplasmic reticulum membrane) substrate cleaved by SPase I (serine endoprotease). The n, h and c stand for N-terminal, hydrophobic and C-terminal region of the SP.

```
1   MGSSHHHHHH SSGLVPRGSH MSDSEVNQEA KPEVKPEVKP ETHINLKVSD GSSEIFFKIK KTTPLRRLME AFAKRQGKEM
81  DSLRFLYDGI RIQADQTPED LDMEDNDIIE AHREQIGGSP LRNRFNEDHD EFSKDDMARE SFDTEEMYNA FLNRRDSSES
161 QLEDHLLSHA KPLYDDFFPK DTSPDDDEDS YWLESRNDDG YDLAKRKRGY DDEEAYDDFD EVDDRADDEG ARDVDESDFE
241 EDDKLPAEEE SKNDMDEETF EDEPEEDKEE AREEFAEDER ADEREDDDAD FDFNDEEDED EVDNKAESDI FTPEDFAGVS
321 DEAMDNFRDD NEEEYADESD DEAEEDSEET ADDFEDDPED ESDETFRDEV EDESEENYQD DTEEGSEIKQ NDETEEQPEK
401 KFDADKEHED APEPLKEKLS DESKARAEDE SDKSEDAAKE IKEPEDAVED FEDGAKVSED EAELLDDEAE LSDDEAELSK
481 DEAEQSSDEA EKSEDKAEKS EDEAELSEDE AKQSEDEAEK AEDAAGKESN DEGKKREDEA VKSKGIARDE SEFAKAKKSN
561 LALKRDENRP LAKGLRESAA HLRDFPSEKK SKDAAQGNIE NELDYFKRNA FADSKDAEPY EFDK
```

**Fig. S26** $His_6$-SUMO-AGARP sequence with acidic residues of AGARP marked in red, and SUMO residues in blue.

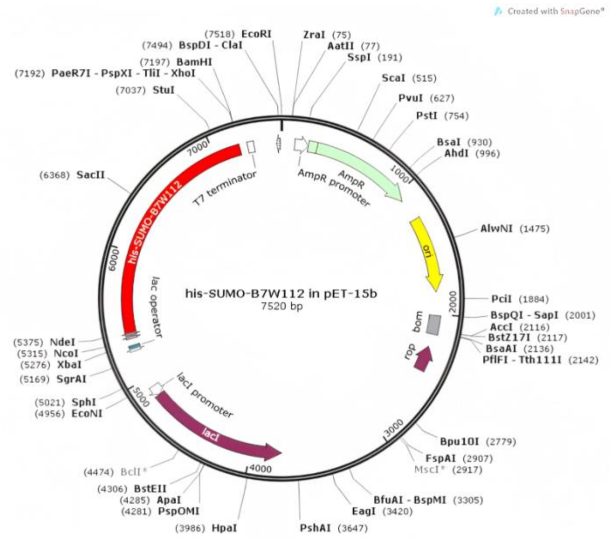

**Fig. S27** pET-15b plasmid with His$_6$ tagged SUMO-AGARP sequence (red) cloned between NdeI and XhoI restriction sites.

We expressed and purified AGARP for the first time. The expression of His$_6$-SUMO tagged AGARP was induced by 0.5 mM IPTG and carried out in 15 °C for 4 hours (**Fig. S26a**). The temperature of bacterial cell culture was set to lower value than the optimum temperature of 37 °C to hinder the heat denaturation of proteins but, most importantly, to suppress the activity of proteases that could enzymatically cleave potentially harmful highly acidic protein.

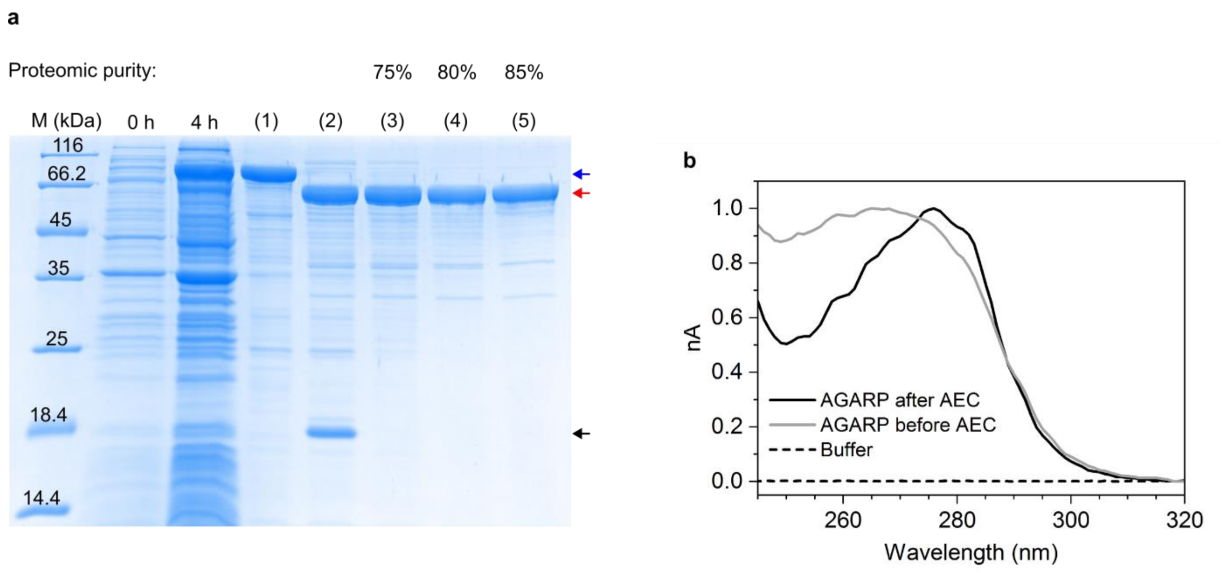

**Fig. S28** (**a**) SDS-PAGE analysis of AGARP expression and purification. 1st band – molecular weight marker; 2nd and 3rd band – cell lysate before induction (0 h) and 4 hours (4 h) after induction of $His_6$-SUMO-AGARP (blue arrow) expression with IPTG at 0.2 mM, respectively; (1) $His_6$-SUMO-AGARP purified on HisTrap column; (2) Cleavage of the $His_6$-SUMO tag (black arrow) from AGARP (red arrow); Subsequent AGARP purification by Ni chromatography (3), anion exchange chromatography (4) and size exclusion chromatography (5). (**b**) Absorption spectra of AGARP before and after anion exchange chromatography (AEC).

Both $His_6$-SUMO-AGARP and AGARP bands are found above the band corresponding to the nominal mass of the protein. This is due to the high negative charge of these proteins, which weakens the interactions with SDS. The molecular masses of the obtained proteins were confirmed by mass spectrometry (**Fig. S27**), 58 330 ± 60 Da for AGARP and 71 630 ± 70 Da for $His_6$-SUMO-AGARP. The mass of AGARP is in agreement with the expected value calculated by ProtParam,[9] 58 338.61 Da, within experimental error. The nominal mass of $His_6$-SUMO-AGARP from ProtParam[9] is 71 745.59 Da, 116 Da less than the experimental result, suggesting that the N-terminal methionine residue (131.2 Da) was cleaved during the protein expression.

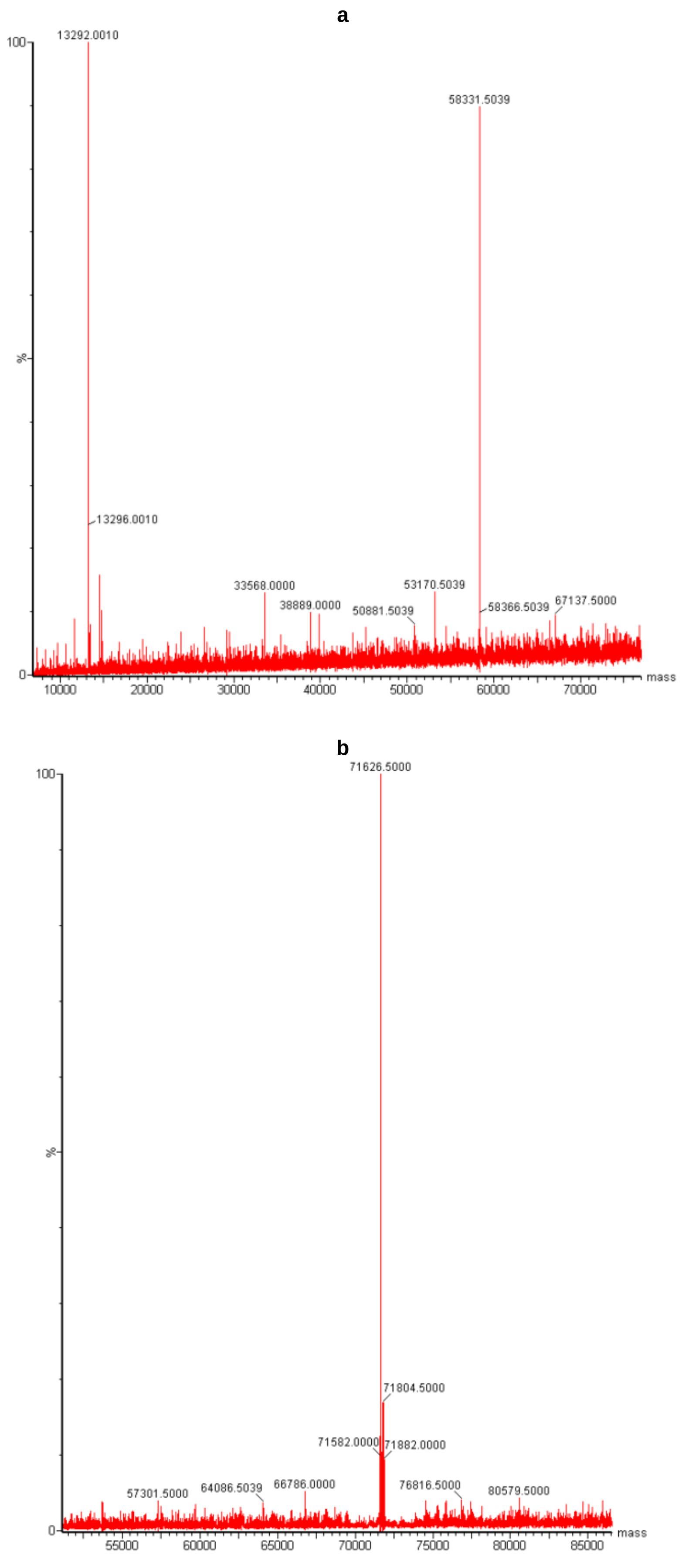

**Fig. S29** Mass spectrum of (**a**) AGARP and (**b**) $His_6$-SUMO-AGARP.

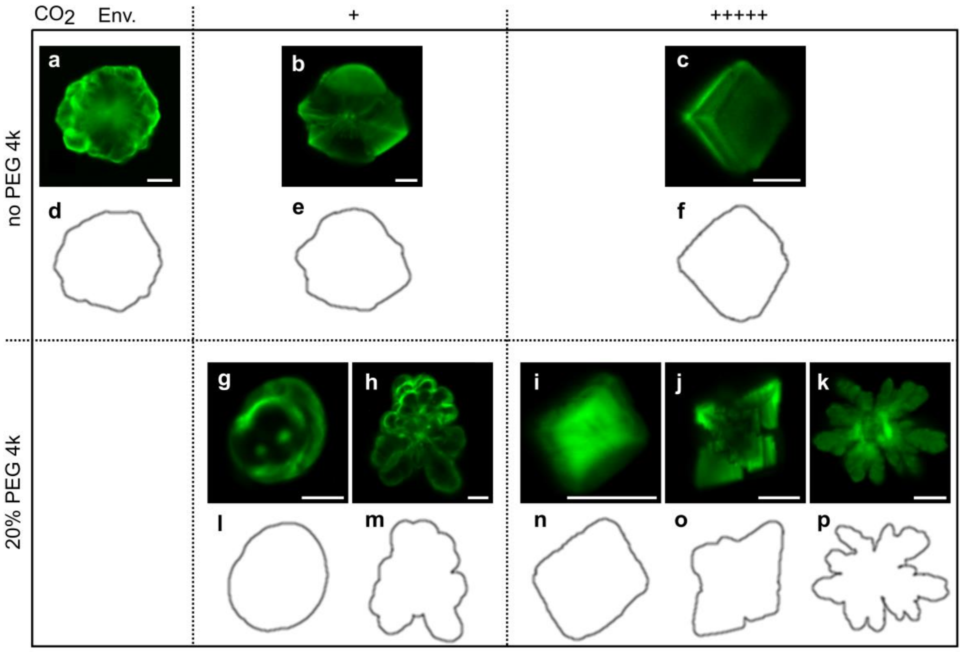

**Fig. S30** (**a**, **b**, **c**, **g**, **h**, **i**, **j**, **k**) Fluorescence confocal images of $CaCO_3$ phases with incorporated AF488-labelled AGARP after rescaling to identical pixel dimensions, resizing to obtain the same percentage of area occupied by the $CaCO_3$ phases and automatic brightness adjustment. (**d**, **e**, **f**, **l**, **m**, **n**, **o**, **p**) Corresponding final images analyzed by FracLac[10]. Scale bar is 10 μm.
